# A new long-read RNA-seq analysis approach identifies and quantifies novel transcripts of very large genes

**DOI:** 10.1101/2020.01.08.898627

**Authors:** Prech Uapinyoying, Jeremy Goecks, Susan M. Knoblach, Karuna Panchapakesan, Carsten G Bonnemann, Terence A. Partridge, Jyoti K Jaiswal, Eric P. Hoffman

## Abstract

RNA-seq is widely used for studying gene expression, but commonly used sequencing platforms produce short reads that only span up to two exon-junctions per read. This makes it difficult to accurately determine the composition and phasing of exons within transcripts. Although long-read sequencing improves this issue, it is not amenable to precise quantitation, which limits its utility for differential expression studies. We used long-read isoform sequencing combined with a novel analysis approach to compare alternative splicing of large, repetitive structural genes in muscles. Analysis of muscle structural genes that produce medium (*Nrap* −5kb), large (nebulin - 22 kb) and very-large (titin - 106 kb) transcripts in cardiac muscle, and fast and slow skeletal muscles identified unannotated exons for each of these ubiquitous muscle genes. This also identified differential exon usage and phasing for these genes between the different muscle types. By mapping the in-phase transcript structures to known annotations, we also identified and quantified previously unannotated transcripts. Results were confirmed by endpoint PCR and Sanger sequencing, which revealed muscle-type specific differential expression of these novel transcripts. The improved transcript identification and quantification demonstrated by our approach removes previous impediments to studies aimed at quantitative differential expression of ultra-long transcripts.

## INTRODUCTION

Cells use alternative splicing to generate a diverse array of protein products from a set of genes(Nilsen and Graveley 2010). Over 90% of genes undergo alternative splicing(Wang et al. 2008). Protein isoforms created from alternative splicing of the same gene may have strikingly different functions and interactions(Yang et al. 2016). Splicing of a gene such as *Mef2D* - a transcription factor required for muscle differentiation, can result in protein products with opposing functions(Sebastian et al. 2013). On the other hand, mutations that cause abnormal splicing are also implicated in genetic disorders, including muscular dystrophies and myopathies(Cummings et al. 2017; Pistoni et al. 2013; Hoffman and Kunkel 1989).

Skeletal muscle has one of the highest rates of tissue-specific alternative splicing(Castle et al. 2008) and genes coding for contractile proteins such as troponin and tropomyosin served as model systems for early mechanistic studies of alternative splicing(Nadal-Ginard et al. 1991). Since then, studies have been performed on splicing of muscle transcription factors, calcium handling genes, muscle-specific mitochondrial splice isoforms and structural proteins such as titin and nebulin(Nakka et al. 2018). However, muscle structural proteins are one of the most challenging to study due to their size and presence of repetitive domains.

Structural proteins are crucial for providing mechanical support within cells and tissues. For example, sarcomeres, the smallest contractile units of muscle, are supported by the largest protein encoded by the mammalian genome, titin (3.3-3.7 MDa). Titin spans half the sarcomere in length, and is coded by transcripts up to ∼106 kilobases long, with 363 exons(Bang et al. 2001). Titin primarily functions as a molecular spring that helps maintain the sarcomere as muscle cells contract(Linke 2017). The elasticity of muscle cells depends on differential splicing of the titin transcript coding a highly repetitive PEVK region, and repeating units of immunoglobulin (Ig), and fibronectin-type-3 domains(Linke 2017).

Nebulin is a large (∼0.9 MDa) structural protein that binds to titin in the Z-disk and extends along the actin thin filament within the sarcomere(Politou et al. 2002; Wang and Wright 1988). It plays roles in assembling, stabilizing and determining the length of the actin thin filaments,(McElhinny et al. 2005) and helps define the width of the Z-disk(Witt et al. 2006). The human nebulin gene has 183 exons that produce transcripts (up to ∼22 kb) that code for multiple conserved 35 aa simple repeats, called nebulin domains(Labeit et al. 1991). In the central region of the protein are 22 super repeats composed of seven-nebulin domains each, one for each turn of the actin molecule(Labeit and Kolmerer 1995). The super repeat region is alternatively spliced during development and in different muscle tissues(Donner et al. 2006; Buck et al. 2010).

A paralog of nebulin is the nebulin related anchoring protein (*Nrap*)(Luo et al. 1997). *Nrap* is the second largest (∼196 kDa) member of the nebulin family of proteins(Luo et al. 1997; Pappas et al. 2011). It is a scaffolding protein, with roles in myofibrillar assembly and organization during muscle development(Luo et al. 1997; Carroll et al. 2001, 2004; Dhume et al. 2006). Its structure is also highly repetitive, containing nebulin domains and super repeats(Luo et al. 1997). It is primarily expressed in intercalated disks of cardiac muscle and myotendinous junctions of skeletal muscle(Carroll and Horowits 2000; Herrera et al. 2000). Alternative splicing of the *Nrap* gene produces tissue specific isoforms (*Nrap*-c and *Nrap*-s for cardiac and skeletal muscle respectively, up to ∼5.5kb)(Mohiddin et al. 2003; Lu et al. 2008).

Recently, Savarese et al. reported the most comprehensive study on alternative splicing of titin by RNA sequencing (RNA-seq) of twelve human muscle types from eleven non-neuromuscular disease patients(Savarese et al. 2018). RNA-seq has been indispensable for studying alternative splicing in whole transcriptomes(Wang et al. 2009). Short-read RNA-seq methods (50-300 bp reads) are an excellent tool for determining whole gene expression and differential exon usage(Trapnell et al. 2010; Li et al. 2015), but the limited read-lengths makes it challenging to determine the full-length and complete splicing patterns of transcripts (Supplementary Figure S1). Further, mapping short-read sequences to highly repetitive regions in the reference genome can be computationally difficult, create ambiguities(Treangen and Salzberg 2011) and reconstructing whole transcripts from short-read data are prone to false positives and artifacts(Steijger et al. 2013). Also, accurate predictions of the complete transcript of a large gene with a vast number of potential exon combinations, is nearly impossible - the 363 exons in titin could theoretically produce over a million alternative splice isoforms(Guo et al. 2010).

Long-read sequencing technologies such as Pacific biosciences (PacBio) isoform sequencing (IsoSeq) are able to sequence full-length transcripts up to 10 kb in length(Wang et al. 2016), with a better error rate than other popular emerging long read sequencing technologies, such as Oxford Nanopore(Buck et al. 2017). However, the entire length of longer transcripts such as titin and nebulin still cannot be captured within maximum IsoSeq read lengths. In addition, IsoSeq is a transcript discovery tool as the bioinformatics pipeline does not support transcript quantification.

Here, we have addressed above deficits of the PacBio long-read IsoSeq technology and used it to compare alternative splicing of transcripts between different murine muscle types: fast skeletal, slow skeletal and heart (Supplementary Figure S2 and Table S1). Our study focuses on the large structural genes: titin, nebulin and *Nrap* to test the limits of IsoSeq. We present a new analysis approach to identify and quantify in-phase transcript structures or consecutive multi-exon stretches of these three transcripts. We have successfully phased exons of known splicing patterns, discovered multiple novel patterns and determined the percent expression of each transcript between tissue subtypes. To our knowledge, this is the first study to apply an exon phasing approach that results in accurate quantification of very long and difficult transcripts. This method improves the utility of IsoSeq, by expanding its potential beyond isoform discovery, to analysis of differential expression.

## RESULTS

### Oligo-dT internal priming provides coverage across large transcripts (> 10kb)

For preliminary analysis between muscle types, the aligned data for all three samples were surveyed in the integrated genomics viewer (IGV). Genes with transcripts close to the 6 kb average read length were often fully covered by a single continuous read (e.g. *Nrap*, Myomesins, Myosins). For larger genes, such as titin (106 kb transcript) and nebulin (22 kb transcript), the entire genomic loci were tiled with reads up to the 5’ end of the gene (Figure 1). This was unexpected given that IsoSeq uses a poly-A capture method. However, the read coverage was not evenly distributed. Dramatic shifts in read depth, visually similar to an upside-down “staircase” pattern were observed when viewing the aligned reads across the gene.

**Figure 1.**
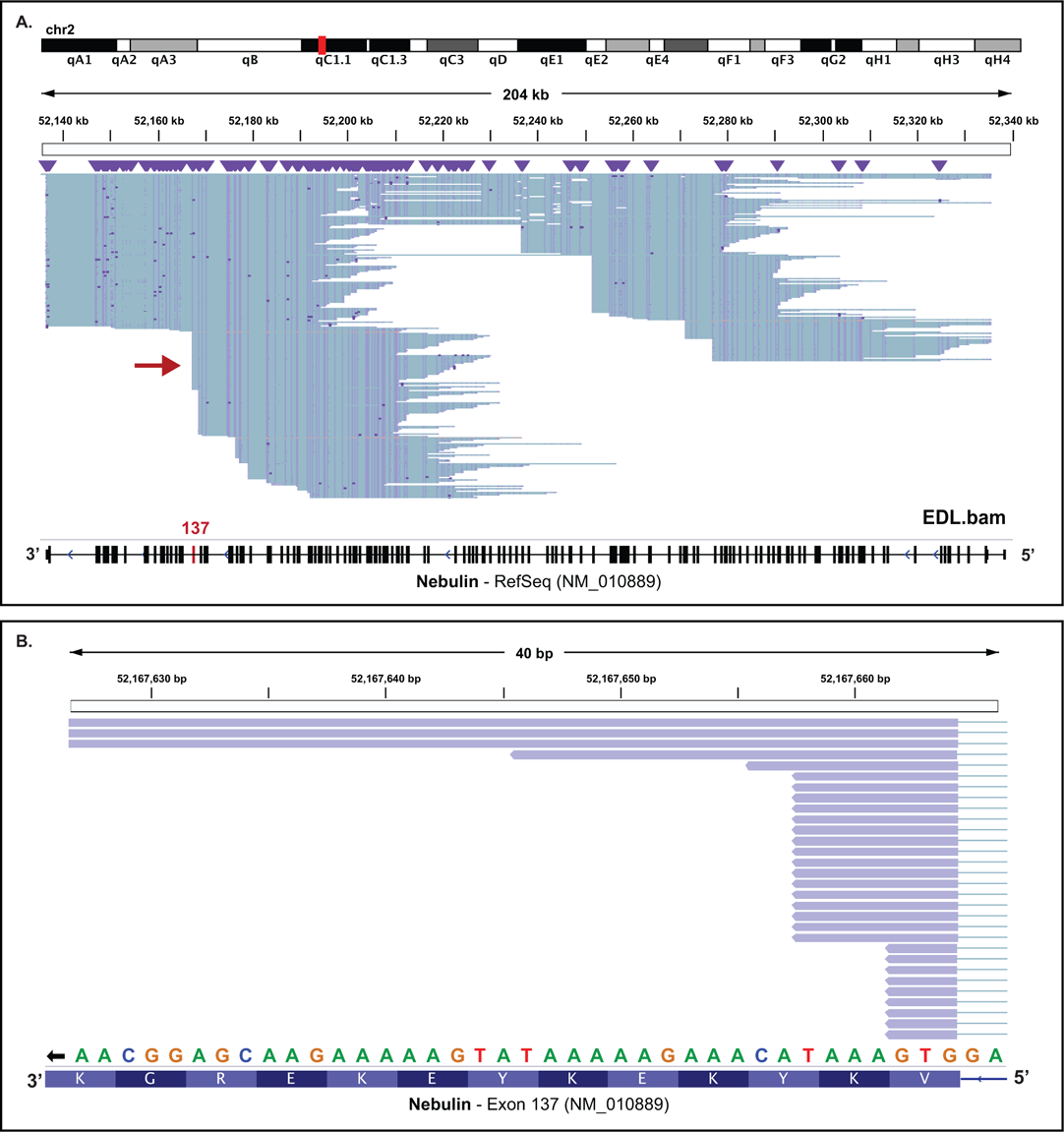
IGV screenshots of reads tiled across mouse nebulin due to internal oligo-dT priming. A) Screenshot of full-length reads (blue/purple lines) tiled across the entire nebulin gene. Nebulin is on the anti-sense DNA strand and is oriented in the right (5’) to left (3’) direction. The mouse nebulin transcript (NM_010889.1) can be up to 22.4kb, or 2x the maximum read length (10kb). Red arrow, the first location upstream of the 3’-end to map several reads due to internal priming. B) Zoomed image of exon 137 showing reads produced from cDNAs generated by internal priming of oligo-dTs. Reference sequence shows consecutive ‘A’ bases followed by non-A bases downstream of where the reads align. Purple lines are coding/exon regions of a reads. Thin connecting blue lines signify intronic sequences that were spliced out. The black lines, dots, and arrowheads are indel errors.

Upon close inspection, reads that aligned upstream of the poly-A region (last 3’ exon) of the gene consistently started on specific exons in all three samples. Focusing in on these exons (e.g. exon 137 of nebulin), the 3’ ends of these reads aligned within 20 bases upstream of multiple stretches of ‘A’ bases intermixed in the genomic sequence (Figure 1B). These reads were likely generated from cDNAs synthesized with internally primed oligo-dT primers. Internal priming has long been described with the use of oligo-dT primers for cDNA generation(Nam et al. 2002). With older technologies such as expressed sequence tags and microarrays, internal priming is viewed as a source of false splice variants. Modern oligo-dT primers used in our study reduce internal priming by adding a random addition of two non-T bases to the 3’ end to anchor them to the polyA tail. However, stretches of ‘A’ bases within an exon followed by non-A bases can circumvent anchoring oligo-dT primers. These observations suggest that coverage of ultra-long transcripts can benefit from internal priming in long-read sequencing if the gene contains enough exons that support it. Other large genes such as Obscurin (27 kb transcript) do not have enough exons that support internal priming for complete coverage (data not shown).

Internal priming also produces reads that do not represent complete mRNA transcripts, creating a source of variability in exon coverage across the length of the gene that may be problematic for exon-based analyses using short-read RNA-Seq.

### Observing splice transcripts across samples

To survey splice differences between tissues, nebulin transcripts were scanned for variable regions. The 3’ region of nebulin is known to be differentially spliced between muscle types, and our long-read sequencing data confirms those reports(Donner et al. 2004). A stretch of exons between 137 and 152 was highly variable (Figure 2), and three novel exons were detected (between exon 147 and 148 of transcript NM_010889.1) that are unannotated in the RefSeq mouse database. Two of these were annotated in the more comprehensive mouse Gencode mouse database (mm10, release M10), but were part of two short nebulin transcripts. One of three exons between 147 and 148 was unannotated (u-002). These three exons are conserved and expressed in 3 out of 4 human nebulin transcripts (RefSeq). Four exons between exon 85 and 86 of NM_010889.1 (or exons 2, 3, 4 and 5 of ENSMUST00000028320.13) were expressed in all Soleus and EDL transcripts and also conserved in human (data not shown). These results suggest that long-read isoform sequencing is able to resolve complex splice isoforms in highly repetitive genes such as nebulin.

**Figure 2.**
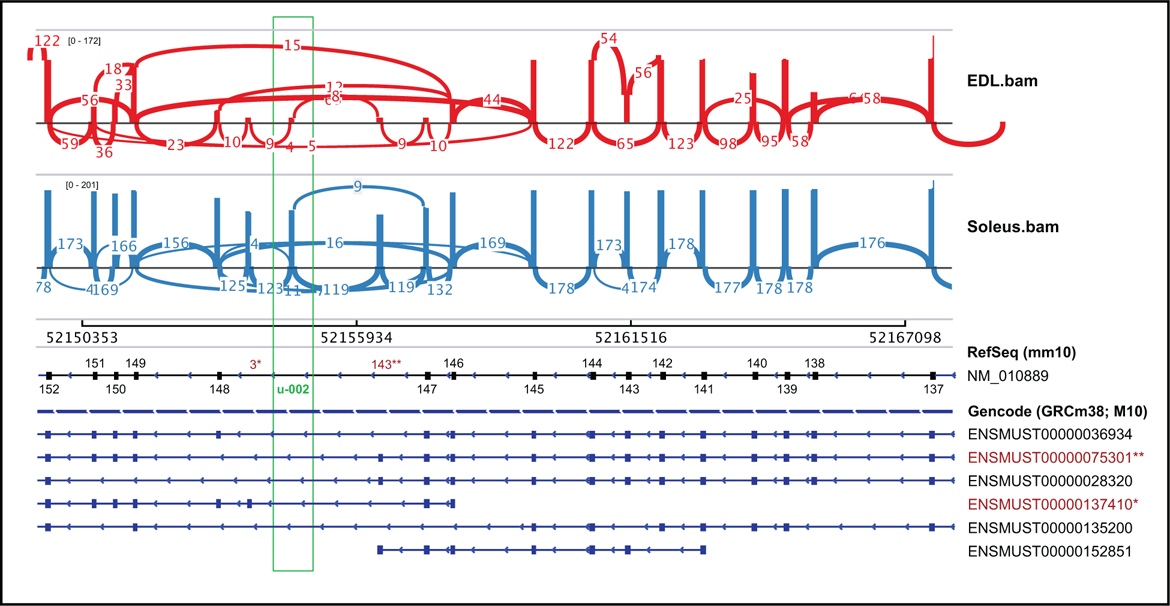
EDL Sashimi plot showing extensive alternative splicing of EDL and Soleus transcripts between exons 152-137 of nebulin. Three exons between exons 147 and 148 are not annotated in the RefSeq mouse database. Two of these exons highlighted in red can be found in the more comprehensive Gencode M10 mouse database. One exon (u-002, green box) is not annotated in either database.

### Identifying unannotated exons and quantifying differential exon usage

To identify unannotated exons and compare differential exon usage between tissues, we first selected for genes with at least 10 cluster reads produced from the IsoSeq method. This threshold resulted in 686 out of 48,440 genes that were adequately covered for further analysis. To reduce the complexity of the splice analysis, we collapsed the gencode (release M10) annotation file using a python script included in the DEXSeq R package, ‘dexseq_prepare_annotation.py’. This produced a collapsed annotation file containing one meta transcript per gene (Supplementary Figure S3A and S3B). Different components of the transcript were renamed as unique exonic parts. See DEXSeq manual for details.

Next, we developed the exCOVator script to perform a preliminary exon-based analysis (exons are treated independently from their neighbors) to determine exon usage between very large transcripts that are not fully covered by the IsoSeq maximum read-length. The script identifies unannotated exons and calculates percent spliced-in (PSI) values for each exon from multiple samples at once using the HT-Seq python library (Supplementary Figure S3C and S3D). Unannotated exons are identified by screening reads for exons missing from the meta transcript of the collapsed annotation file. This identified 1,218 unannotated exons that were added into the meta transcripts of the collapsed gencode annotation file. Using this new collapsed annotation file, sample cluster reads that matched the annotated exonic parts were counted. The full-length read counts from each cluster read was extracted and used for PSI calculations for each exonic part. Finally, the full-length PSI was compared across samples. We selected for exonic parts that were at least 20% difference in PSI between at least two out of three samples to reduce artifactual differences. This resulted in 2,631 unique exonic parts differentially spliced in 433 genes between at least two of the three muscle samples.

For this study, we focused only on analyzing three large muscle structural genes: 1) *Nrap*, that produces ∼6kb transcripts which fall within average read-lengths, 2) nebulin, that produces ∼22kb transcripts that are over twice as long as the maximum read length and 3) titin, that produces 106kb transcripts that are over 10 times the read-length.

### Novel muscle type specific splicing patterns in nebulin related anchoring protein

Differential usage of exon 12 of *Nrap* was observed between all three muscle tissues. Lu et al. described the expression and alternate splicing of *Nrap* in skeletal and cardiac muscle during development(Lu et al. 2008), and identified two splice variants: 1) *Nrap*-s, a splice transcript of *Nrap* that includes exon 12 (relative to RefSeq annotation NM_008733.4) and is exclusively expressed in skeletal muscle and 2) *Nrap*-c, a splice transcript that is missing exon 12 and is exclusively expressed in cardiac muscle. The present analysis confirms their findings that there is no expression of exon 12 in *Nrap* transcripts in cardiac tissue (0/567 reads). In contrast to their findings, our data suggests that *Nrap*-c is not exclusively expressed in cardiac tissue. Both *Nrap*-s and *Nrap*-c splice isoforms are expressed in skeletal muscle and there is a 53% higher usage of exon 12 among the transcripts in the predominantly fast-twitch EDL (576/862 reads; 67% PSI) compared to the slow-twitch soleus muscle (377/2621 reads; 14% PSI; shown as exonic part 38; Figure 3A and 3B).

**Figure 3.**
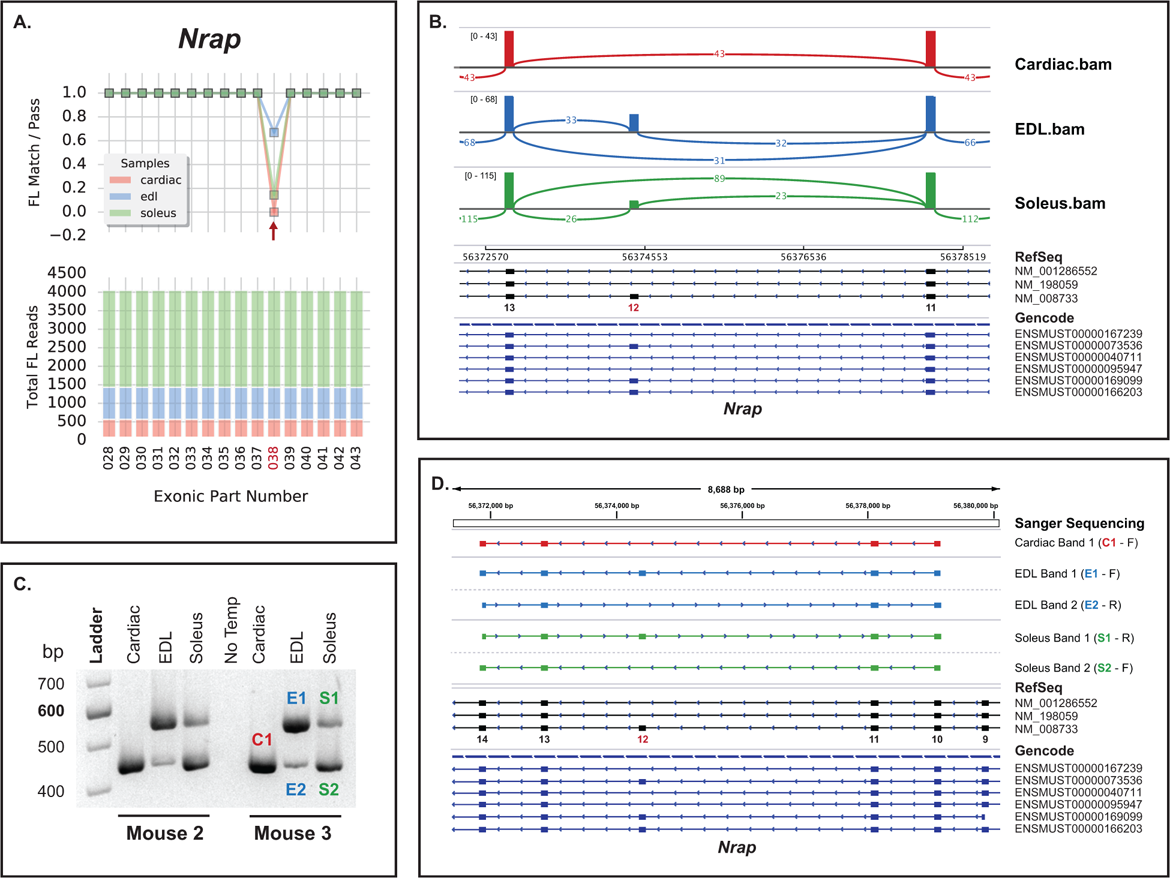
*Nrap* has differential usage of exon 12 (exonic part 038) in all three mouse muscles. A) *Nrap* gene graph (cropped) produced by ‘exCOVator.py’. Bottom stacked bar graph showing total full-length (FL) read coverage of all exonic parts (EP) for the gene. Top line graph, displaying the ratio of FL reads matching EP sequence / total FL reads that cover the EP coordinates. B) Sashimi plot exported from the integrated genomics viewer (IGV) displaying differential splicing of exon 12. C) Agarose gel showing RT-PCR validation products generated from primers that target exons 9-14 (549 bp; includes exon 12 and 444 bp excludes exon 12). The gel shows differential exon usage between soleus, EDL and heart of two independent mice. NT= no transcript control. D) Sanger sequencing of products excised from gel in panel C. Top half shows sequences aligned using the IGV BLAT tool. Cardiac band 1, and EDL and soleus band 2 are missing exon 12.

To validate the findings, we performed RT-PCR and Sanger Sequencing using the same primers in the publication(Supplementary Table 2)(Lu et al. 2008), and total RNA extracted from muscles of two independent mice (Figure 3). Our RT-PCR results confirm the IsoSeq analysis data, showing: 1) low usage of exon 12 in soleus transcripts (more 444 bp product than 549 bp), 2) high usage of exon 12 in EDL transcripts (more 549 bp product than 444bp) and 3) no usage of exon 12 in cardiac transcripts (single 444 bp product). Sanger sequencing of the excised bands further confirmed the splice products (Figure 3D). By corroborating the findings of the prior study, the present results establish that PacBio long-read IsoSeq can be used as a semi-quantitative analysis approach for differential exon usage between samples. Additionally, the IsoSeq data extends the prior data and show differential exon usage of exon 12 among different skeletal muscle types, as both *Nrap*-s and *Nrap*-c transcripts are expressed in fast and slow skeletal muscle at varying amounts. Not only is there alternative splicing of *Nrap* transcripts between cardiac, and skeletal muscles, but also within skeletal muscle types. This suggests that *Nrap*-s and *Nrap*-c are misnomers.

### Nebulin exon 138 is differentially used between slow and fast skeletal muscle

Our preliminary observations and exCOVator found multiple differentially used exons between 137 and 152 of nebulin (NM_010889.1; Figure 2, 4A and 4B). We used exon 138 to validate differential exon usage in this region (Figure 4C and 4D). The results showed a single RT-PCR product at ∼500 bp in soleus muscles, suggesting that exon 138 is present and that there is a consistent retention of all exons between 135 and 139 in the majority of transcripts in the soleus muscle (Figure 4C). The EDL muscles had two product sizes, a ∼500 bp product similar to the soleus and a smaller ∼400 bp product missing exon 138. These results suggest differential exon usage of 138 in skeletal muscles which is excluded from a subset of EDL transcripts, but not in the soleus.

**Figure 4.**
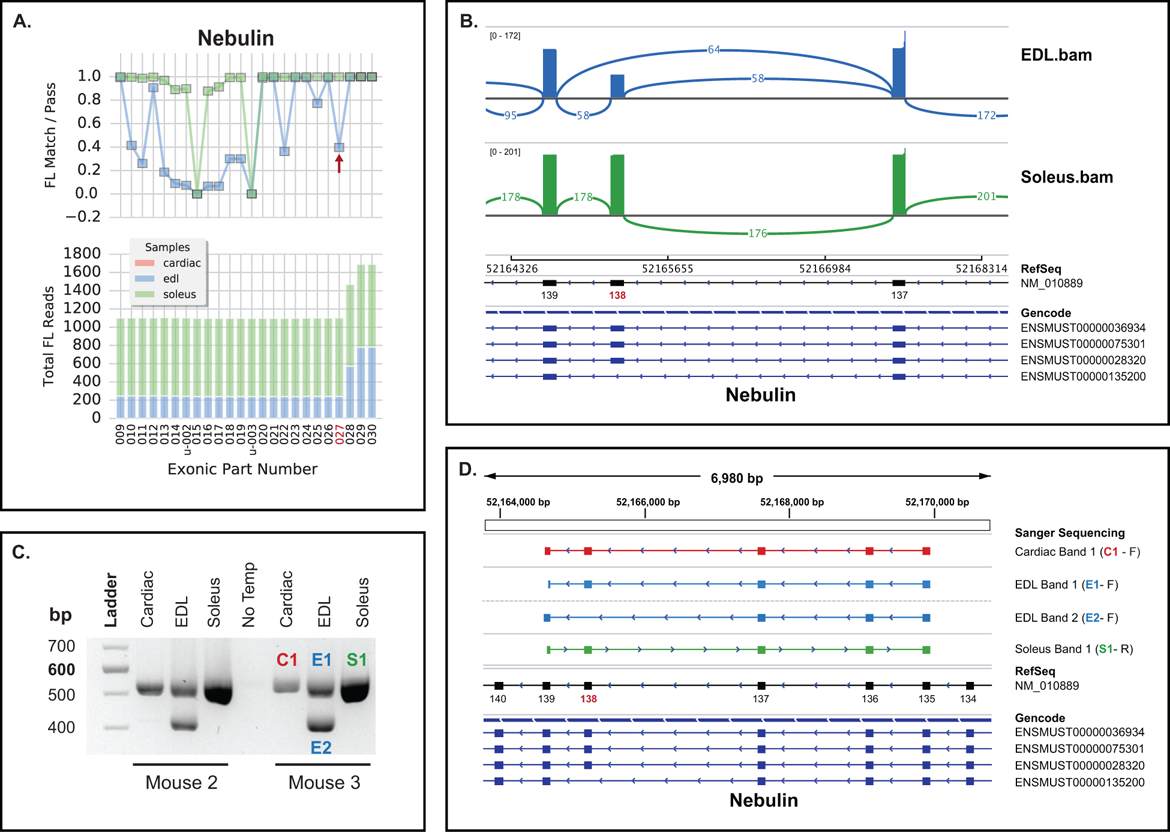
Differential usage of exon 138 of mouse nebulin (NM_010889). A) Nebulin gene graph (cropped) produced by ‘exCOVator.py’. Bottom stacked bar graph displays full-length (FL) read coverage of all exonic parts (EP) for the gene. Top line graph displays the ratio of FL reads that match the EP sequence / total FL reads that cover the EP coordinates. B) Sashimi plot exported from the integrated genomics viewer (IGV) displaying differential splicing of exon 138. C) Agarose gel showing RT-PCR validation products of nebulin exon 138. Shows differential exon usage between soleus, EDL and heart. NT= no transcript control. Cardiac shows similar banding pattern as Soleus, however few reads were detected during sequencing. Primers were designed to target exon 135 and 139. D) Sanger sequencing of products cut from agarose gel seen in panel C. Top portion shows sequences aligned in IGV using the BLAT tool. Only the EDL band 2 (E2 from panel C) is missing exon 138 sequence.

Cardiac muscle has low expression of nebulin(Bang and Chen 2015). However, we identified a prominent single ∼500 bp product similar in size to the product in the soleus (Figure 4C). This product could be the result of the higher sensitivity of RT-PCR compared to IsoSeq. However, the soleus band is overloaded making size determination more ambiguous. It is possible that some sensitivity was lost by selecting for cDNAs > 5kb prior to IsoSeq. Sanger sequencing of the amplicon products show that the cardiac splice product is identical to what is found in the EDL and soleus (Figure 4D), which rules out possible mis-priming to nebulette, a nebulin gene family member that is highly expressed in the heart.

### Titin, cardiac specific unannotated cassette exon 191

The IsoSeq analysis results shows that titin exonic part 129 or exon 191 (NM_011652.3) is almost always used in transcripts of soleus (34/35 FL reads; 97% PSI) and EDL (357/361 FL reads; 99% PSI) muscles, but is missing from 64% of the transcripts in cardiac muscle (84/232 FL reads; 36% PSI; Figure 5A and 5B). Both Gencode and RefSeq databases show this exon to be constitutively expressed. Therefore, the present data suggest titin exon 191 may be an unannotated cassette exon that is specifically removed from a subset of cardiac muscle transcripts. RT-PCR data shows that exon 191 is retained in Soleus and EDL transcripts by the presence of a single product matching the predicted size of 852 bp (Figure 5C). However, the cardiac sample shows three different RT-PCR products. The 615 bp product is predicted to only be missing exon 191. The 882 bp product contains all exons between 190 and 194. The product in between at ∼780 bp is an unknown splice product. Attempted Sanger sequencing of the unknown product resulted in truncated sequences and remains ambiguous (Figure 5D; Cardiac Band 2). This is likely because the product is a heteroduplex between the smaller 615 bp and larger 882 bp splice products. Overall, these data confirm that titin exon 191 is an unannotated cassette exon that is only removed from a subset of mouse cardiac transcripts.

**Figure 5.**
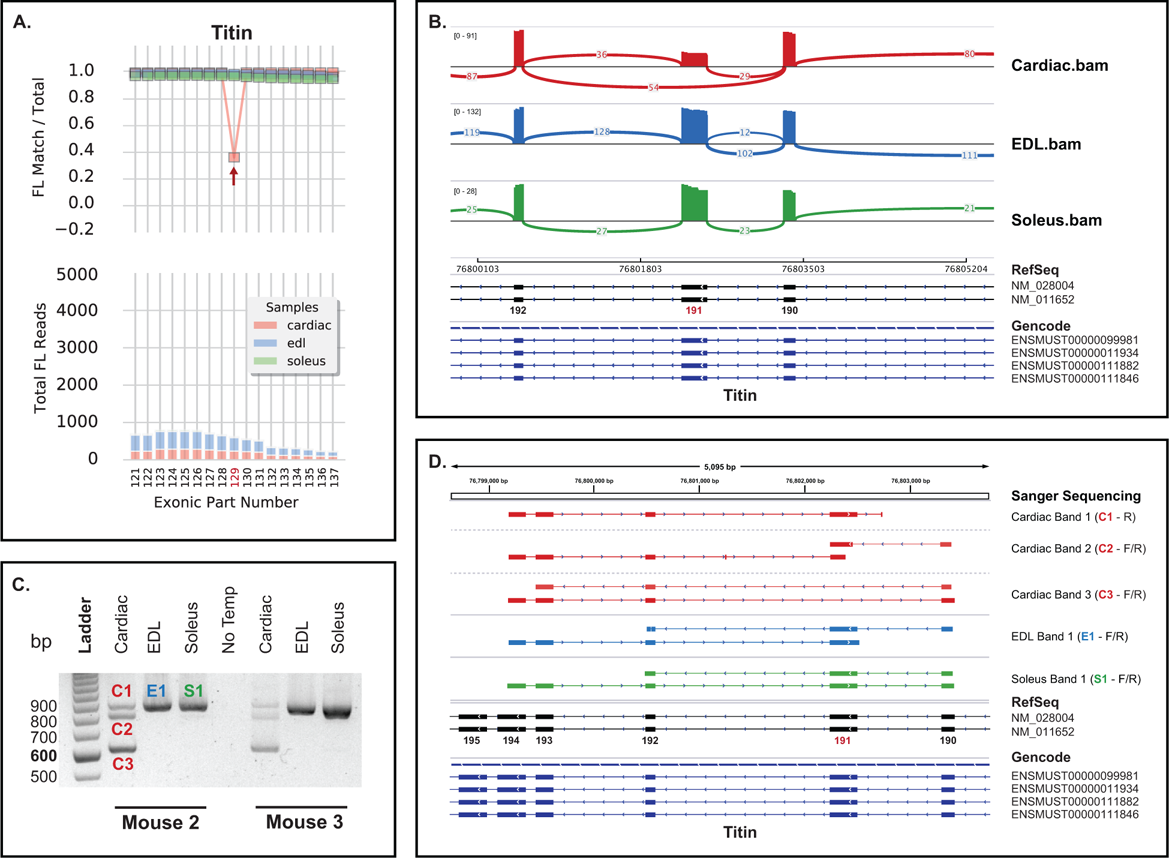
Titin exon 191, found to be an undocumented cassette exon that is spliced out in a subset of mouse cardiac transcripts. A) Exon coverage graph of titin gene (cropped). Top line graph, red arrow points to exonic part (EP) 129 or exon 191 of NM_011652.3 (N2-A). Bottom stacked bar graph, red arrow points to coverage of EP 129 in all three tissues. B) Sashimi plot showing the same data in the integrated genomics viewer (IGV). C) Agarose gel / RT-PCR validation. Primers target exon 189-194 with predicted product sizes of 882 bp (with exon 191, C1) and 615 bp (without exon 191, C3). Soleus and EDL transcripts all include exon 191. Cardiac has both predicted bands and one extra unknown band in the middle (C2) that is likely a heteroduplex. NT = no template control. Primers target exon 190 and 194. D) Sanger sequencing data of bands from figure 5C. Exon 191 is missing only from cardiac band 3 (C3) and is present in all other tissues.

### Annotation and quantification of transcript structures

One of the strengths of long-read sequencing is the ability to phase multiple neighboring exons to determine transcript splice patterns within the same read. However, the exon-based approach used here to identify and quantify differentially used exons does not account for phasing. To address this, we developed another tool, exPhaser, to quantify and annotate splicing patterns of larger transcript structures. The exPhaser script takes in a bed file of exons and sample files (bams) and determines the splicing pattern of input exons within each sample read. Then it outputs a table of FL-read counts for each splicing pattern, and all transcript annotations that match the pattern. As exon input for exPhaser, cassette exons were selected that defined the isoform according to known annotations and exCOVator output (e.g. novel exons and differentially used exons). We also filtered out unannotated isoforms with < 10 FL-reads or after visually determining in IGV that they were read/alignment errors.

### Phasing of full-length *Nrap* transcripts

Most reads aligned to *Nrap* span the entire length of the transcripts, therefore we annotated and counted all transcripts by phasing all known cassette exons: 2, 12, 17, 37, 38, 39, and 40 (NM_008733; Figure 6A). The differential expression of each transcript reflected the results of the exCOVator output (Figure 3 and 6A). The primary transcript expressed in cardiac tissue is ENSMUST00000095947 (excludes only exon 12) at 97.5% (541/555 FLreads). In the EDL muscle, ENSMUST00000040711 is expressed at 33.5% (280/837 FL reads) and ENSMUST00000073536 (all exons) at 66.5% (557/837 FL reads). The soleus expresses more of ENSMUST00000040711 at 85.6% (2,177/2,544 FL reads) and ENSMUST00000073536 at 13.2% (335/2,544 FL reads). Additionally, we identified a rare transcript ENSMUST00000095947 (excludes exon 2 and 12) exclusively expressed in cardiac at 2.5% (14/555 FL reads) and soleus muscle at 1% (26/2544 FL reads).

**Figure 6.**
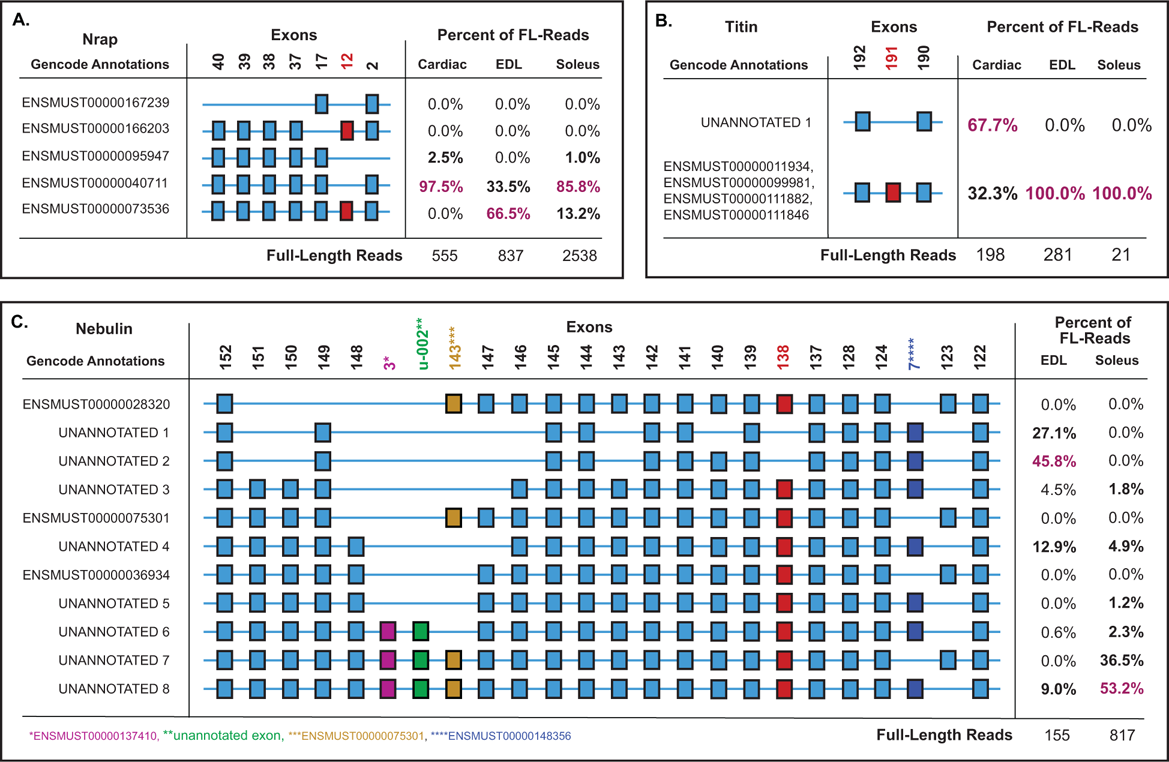
Quantifying mouse *Nrap*, titin and nebulin transcript isoforms that include a validated exon in different muscles using exPhaser. A) NRAP transcript structures with exon numbering based on NM_008733. B) Titin transcript structures with exon numbering based on NM_011652. Some splice patterns could not be uniquely mapped to a single annotation. C) Nebulin transcript structures with exon numbering based on NM_010889. Blue squares are isoform defining exons. Red squares are in red were validated using RT-PCR and Sanger Sequencing.

### Phasing nebulin exons in the Z-disk and 3’ end of the super repeat region

After analyzing differential exon usage using exCOVator, the results showed that differential splicing of nebulin only occurred in the 3’ half of the transcript. We also surveyed other regions upstream known to be alternatively spliced such as exons 65-67 and 83-87. No differential splicing was detected between EDL and Soleus in these exons.

Differential splicing of the 3’ end of nebulin transcripts occurred in 14 exons between 122-152 of NM_011652 (Figure 2 and 6C). These exons code for nebulin peptides found within the Z-disk and 3’ end of the super repeat region. This includes two mutually exclusive exons located between exons 122-124, and twelve cassette exons between exons 137-152. After phasing all 14 differentially spliced exons and additional flanking exons, we found all transcripts structures expressed in the three muscles were not annotated in RefSeq or Gencode (M10) databases. This was because most reads contained a combination of one or more of the exons from other annotated transcripts or the unannotated exon. Corroborating our exon-based analysis, the transcripts expressed in the EDL have more exons spliced out between exons 137-152 (Figure 4A). This includes differential splicing of exons 138, 140, 143, 146, 147, 143*** (of ENSMUST00000075301), u-002, 3* (of ENSMUST00000137410), 148, 150, and 151. Only 9% (14/155 FL reads) of EDL reads retain all exons between 137 and 152. In contrast, the Soleus retained all of the listed exons in 89.6% (733/817 FL reads) of transcripts. It is known that the width of the sarcomeric Z-disk in fast-twitch muscle (e.g. EDL) is narrower compared to slow-twitch muscle (e.g. Soleus). Alternative splicing of the Z-disk region of nebulin transcripts of EDL and Soleus correlate with Z-disk width observations.

In addition, we analyzed the splicing pattern of two mutually exclusive nebulin exons referred in the literature as exon 127^♦^ and 128^♦^(Figure 6C)(Donner et al. 2006). Translated to our exon numbering using NM_010889, exon 127^♦^= 123 and exon 128^♦^is not annotated in RefSeq mm10, but instead annotated as exon 7**** of ENSMUST00000148356 in the mouse Gencode database. Our phasing results show that transcript structures in the EDL all include exon 7**** (155/155 FL reads). On the other hand, the Soleus expresses a mixture of transcripts that contain exon 7**** and exon 123, but with slightly more transcripts expressing exon 7**** at 63% (514/817 FL-reads) than exon 123 at 37% (303/817 FL reads) which agrees with previous studies(Donner et al. 2006).

Based on RefSeq annotation NM_001271208, alternative splicing of human nebulin occurs in exons 63-66 (66-69 of NM_010889 in mouse), 82-105 (83-87), 143-144 (124-7**** of ENSMUST00000148356), and 169-173 (4 exons between 148-149 and 149 itself)(Donner et al. 2004, 2006; Lam et al. 2018). We found alternative splicing of all of these exons to be conserved in mouse except for exons 63-66 (data not shown) and 82-105 (Supplementary Figure S4) that are constitutively expressed.

### Phasing multiple transcript structures of titin

For titin, we could not phase all differentially spliced exons at once because they are too far apart to be contained within the length of a single read. To illustrate this, we compared the expression of titin N2-A (NM_011652, principle skeletal isoform) vs N2-B (*NM_028004, principle cardiac isoform) transcripts between the three muscle types, using the N2-A isoform for exon numbering unless noted. The N2-B transcript excludes exons 47-167 of N2-A, but includes an alternate exon 45*. Initially, we sought to phase transcripts using exon 45* (N2-B specific), 46, 47 (N2-A specific) and 168. However, the distance between exon 45* and 168 in N2-A transcripts is much greater than the 10kb maximum read length. Therefore, we phased the skeletal and cardiac specific exons using two groups of exons (Figure 7A and 7B). In the skeletal muscle isoform analysis, we phased exons 45*, 46, 47, 48, 49, and 50. Results confirmed that N2-A is the primary isoform expressed in skeletal muscle (Soleus: 19/19 FL reads and EDL: 160/160 FL reads) and that cardiac muscle did not express exons 47-50 (202/202 FL reads; Figure 7A). In the parallel cardiac isoform analysis, we phased exons 45* 46, 47, 167, 168, and 169. Those data showed all transcripts expressed in cardiac muscle were missing exons 47 and 167, but included exons 45*, 46, 168, and 169 aligning with titin isoform N2-B (Cardiac: 190/190, EDL: 0/0, and Soleus: 0/0 FL reads).

**Figure 7.**
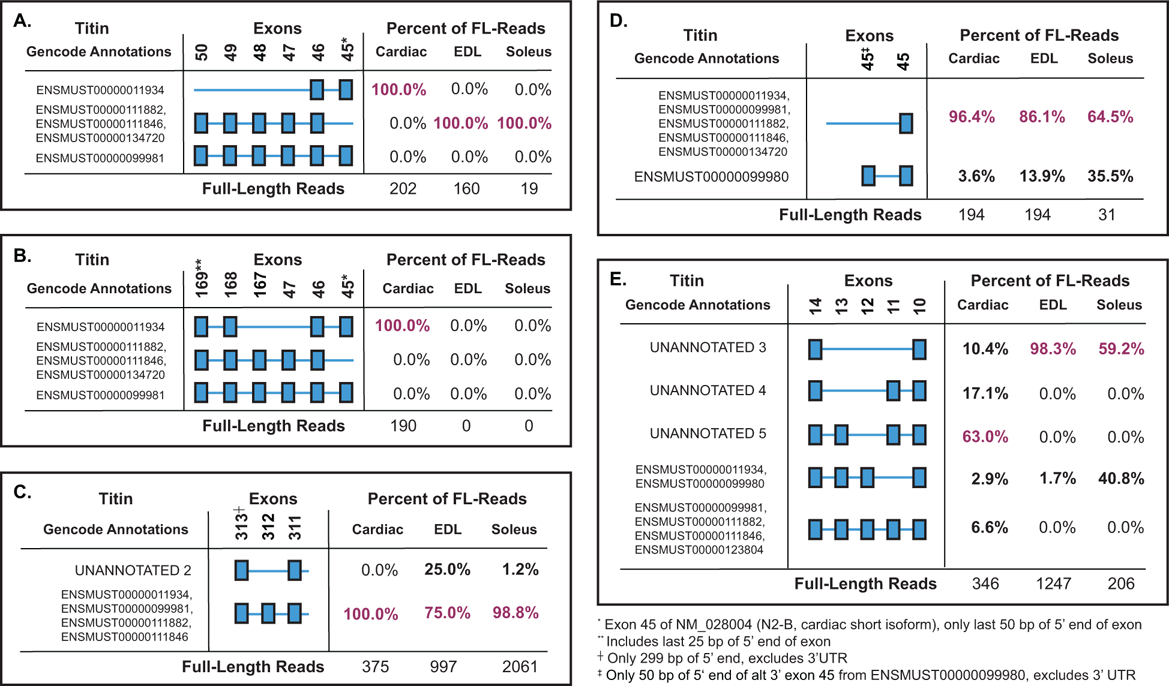
Phasing of additional titin exons and quantification of transcript structures using exPhaser. Exons are numbered by NM_011652/ENSMUST00000111846 (N2-A, principle skeletal long isoform) unless noted. A) Quantifying transcript structures focusing on N2-A isoform defining exons (47, 48, 49, 50) and some N2-B a cardiac defining exon (45*). B) Quantifying transcript structures focusing on N2-B (cardiac) isoform defining exons (45, 46,168, 169) and two N2-A defining exons (47 and 167). C) Exon 312 of titin spliced out in skeletal muscle transcripts, an unannotated skeletal muscle specific isoform. D) Phasing of exon 45‡ of ENSMUST00000099980 an alternate 3’ terminal exon that is differentially used in all three tissues. E) Phasing of exons 11,12 and 13 showing that the most abundant isoforms in each tissue are unannotated. Transcript structures listed with multiple ID’s cannot not be uniquely annotated using the exons provided.

Next, we phased four other sets of differentially used exons in titin including exon 191, exon 312, exon 45^‡^ of ENSMUST00000099980 and exons 11, 12 and 13 (Figure 7E). First, we phased exon 191, the validated cassette exon that is only removed from a subset of cardiac muscle transcripts (Figure 6B). The cardiac phasing data were similar to the RT-PCR results (Figure 5) with 32.3% (64/198 FL reads) of transcripts having exon 191 spliced-in. In both skeletal muscles, exon 191 is found in all transcripts (EDL: 281/281, Soleus: 21/21 FL reads) suggesting it is a cassette exon that is only removed from cardiac muscle transcripts. Further, mouse exon 191 is in-frame and conserved in human as exon 191/312 of NM_133378 or as exon 243 based on the Locus Reference Genomic (LRG) coordinates used for clinical reporting. To determine if alternative splicing of exon 191 is also conserved in humans, we consulted the Cardiodb website for titin expression in human heart, and the comprehensive short-read RNA-Seq study by Savarese et al on 11 different human skeletal muscle types(Savarese et al. 2018; Roberts et al. 2015). In heart, 59% of exon 191 is spliced-in in left ventricle tissue of dilated cardiomyopathy (DCM) patients (84 end-stage patients), and 69% in the GTEx data set (left ventricle tissue, 105 samples). The two exons flanking 191 are almost constitutively expressed with a spliced-in rate of 96-100% in DCM patients and GTEx. Savarese et al. show that exon 191 (LRG exon 243) is constitutively expressed across 11 adult skeletal muscle types. These data corroborate our findings that exon 191 is a cassette exon that is only spliced out from a subset of cardiac and not from skeletal muscle transcripts. However, it also suggests that there are slight differences in splicing of exon 191 in mouse compared to human. These discrepancies could be due to differences in tissue sampling locations. The human heart studies used the left ventricle in their sequencing experiments. In comparison, we used the apex of the mouse heart which includes the lower portions of the left and right ventricles.

Second, we phased the penultimate exon of titin, exon 312, along with adjacent neighbors. The results showed that exon 312 is always included in cardiac transcripts (375/375 FL reads; Figure 7C). In contrast, EDL and Soleus muscles splice out exon 312 in 25% (249/997 FL reads) and 1.2% (25/2061 FL reads) of transcripts respectively. The phasing and exon-based analysis (data not shown) suggests that exon 312 of titin is a cassette exon that is spliced out specifically from skeletal muscle transcripts. It is also preferentially spliced out of transcripts in the fast EDL compared to the slow Soleus muscle. According to Cardiodb and GTEx, exon 312 (LRG exon 363) is 99% spliced-in in human left ventricle tissue. Data from Savarese et al. show that exon 312 is 91% spliced-in across different skeletal muscles. Additionally, the exon is in-frame in both species. These data support that alternative splicing of exon 312 is conserved and that it is a cassette exon that is only removed from a subset of skeletal muscle transcripts.

There are some notable differences between exon 312 in mouse and humans that do not affect the conclusions. The human equivalent of exon 312 (LRG exon 363) is larger with an additional 7 bp on 3’ end and 53 bp on the 5’end of the exon. However, the extended human exon 312 remains in-frame because its flanking exons are also different lengths and accommodate for the discrepancy.

Third, we phased the alternative 3’ exon 45^‡^ of ENSMUST00000099980. After manually inspecting the reads in IGV, we noted numerous partial reads of the large exon 45^‡^ (data not shown). Therefore, we decided to phase the exon (only 50 bp of 5’ end) and its upstream neighbor, exon 45 of NM_011652. The phasing results show that exon 45^‡^ is expressed in 3.6% (7/27 FL reads) of all cardiac muscle transcripts, 13.9% (27/194 FL reads) of EDL transcripts and 35.5% (97/118 FL reads) of Soleus transcripts. Our data suggests that ENSMUST00000099980 is a relatively minor isoform expressed in all three muscles but is expressed higher in skeletal compared to cardiac muscle.

In humans, the paralog of ENSMUST00000099980 in mouse is the novex-3 (ENST00000360870/NM_133379), the minor small cardiac isoform of titin. The alternative 3’ exon is conserved in humans as exon 46 of NM_133379 (48 LRG) and has a 22% sequence difference compared to exon 45^‡^ in the mouse paralog. According to Cardiodb, exon 46 is expressed in 9.4% and 7.3% of human left ventricle transcripts in the DCM and GTEx respectively. The GTEx data was calculated by taking the ratio of the alternative 3’ exon vs the primary end exon (364 LRG). Their result is close to our 18% expression of exon 45^‡^ found in mouse cardiac tissue. On the other hand, Savarese et al. describe exon 46 as only 2% spliced-in in adult skeletal muscles compared to our 13.9% and 35.5% in mouse EDL and Soleus transcripts, respectively. However, this discrepancy may be due to a combination of species differences, location of cardiac tissue sampled and the poor reliability of percent spliced-in (PSI) calculations on 3’ exons using short-read data.

Lastly, we phased exons 11, 12 and 13 of titin (and their neighboring exons 10 and 14) and observed that the most highly expressed transcripts in each of the three tissues have not previously been annotated (Figure 7E). Our results show that there are extensive differences in the usage of exons 11, 12 and 13 in the three muscle types. Exon 11 is not used in any titin transcripts expressed in the two skeletal muscles and is likely that the N2-A annotation may be incorrect. In contrast, cardiac muscle makes extensive use of exon 11, which appears in 86.7% of expressed transcripts. Exons 12 and 13 seem to be co-expressed within skeletal muscle, but only in less than half of Soleus (40.8%) and very rarely in EDL (1.7%) transcripts. Exon 12 is rarely used in heart (9.5%), but exon 13 is expressed in more than half (69.6%) of expressed transcripts. This suggests that peptides coded by all three exons are dispensable for titin’s function in fast skeletal muscle since 98.3% of transcripts in EDL do not express these exons. There is also similar expression of exon 13 in slow skeletal and cardiac muscle. However, slow skeletal muscle includes exon 12 in more transcripts than cardiac muscle with the inverse expression pattern seen for exon 11.

All three exons are conserved in humans and retain the same numbering. Savarese et al show that exon 11 is constitutively spliced out in human which agrees with our mouse data. They also show exon 12 to have an average of 54% inclusion and exon 13 with a 79% inclusion in skeletal muscle transcripts. Our soleus muscle data agrees with theirs, showing a similar exon inclusion rate of 40.8%. However, in the EDL muscle, exon 12 is only found in 1.7% of transcripts which is a large discrepancy. Our mouse exon 13 results also conflict with their human data. We found exon 13 to be co-expressed in transcripts with exon 12, sharing the same inclusion rate of 1.7% in EDL is vastly lower than their average of 79% inclusion in skeletal muscle. It’s likely that these discrepancies are due to the averaging of values across multiple skeletal muscles in the Savarese et al study.

Exon 11 is expressed in left ventricle tissue at 59% in DCM and 69% in GTEx according to Cardiodb. This agrees with our analysis that cardiac tissue includes exon 11 in 86.7% of transcripts, the highest inclusion rate out of our three muscle tissues. The database reports that exon 12 is included in 79% of left ventricle transcripts in the DCM study and 75% in GTEx. These inclusion rates of exon 12 are much higher than our study in which the exon is included in only 9.5% of mouse cardiac transcripts. Exon 13 is expressed in 96% of DCM and 95% of GTEx transcripts, but only 72.5% in mouse cardiac transcripts. We see some agreement between species such as exon 11 being included exclusively in transcripts expressed in heart, but the other two exons seem to be included at higher rates in humans. These differences could also be due to species specific splicing or different sampling locations of the heart.

## DISCUSSION

Our work here has developed a unique approach to identify and quantify novel isoforms of ultra-long transcripts between samples using long-read sequencing and the IsoSeq approach. On its own, IsoSeq has so far only been used for isoform discovery purposes. Use of our exon-based (exCOVator) approach enables quantifying differential exon usage and identifying novel exons in large structural protein transcripts. We showed this for three such proteins of specific importance in muscle: titin, nebulin and *Nrap*, without resorting to additional short-read data.

This reduces cost, but more importantly, we specifically show that this method is useful for transcripts with highly complex and repetitive sequences, which are difficult to resolve using short-read sequencing and other molecular assays. Using our exon phasing (exPhaser) approach, we determined splicing patterns between cassette exons to generate transcript structures. The exon pattern of these structures was mapped to known annotations for identification and used to count reads to quantify their expression in each sample.

*Nrap* transcript sizes (∼5kb) are within our 6kb average read length making them an ideal test sample for our study. There are also fewer known splice isoforms, with one described as unique to cardiac (*Nrap*-c) and another to skeletal muscle (*Nrap*-s)(Mohiddin et al. 2003; Lu et al. 2008). These isoforms were validated by our exon-based analysis. We found that *Nrap*-c (exon 12 excluded) was indeed the only isoform expressed in heart (Figure 3). We also confirmed that *Nrap*-c is not exclusively expressed in cardiac muscle, but is differentially expressed in skeletal muscles as well, with higher expression of *Nrap*-c in the slow-twitch Soleus than in the fast-twitch EDL muscle (Figure 3C). *Nrap*-s (includes exon 12) was also differentially expressed in the two skeletal muscle types with an inverse pattern to *Nrap*-c. This muscle type difference went undetected in previous studies using a mixture of lower limb muscles(Lu et al. 2008). By phasing every known cassette exon in *Nrap*, we quantified all known *Nrap* isoforms in the three muscle types. Doing so also increased our sensitivity for rare transcript isoforms, leading to the detection of an isoform exclusively expressed in heart and soleus muscles (Figure 6A). This rare transcript lacks both exon 12 and exon 2, the latter being one of two exons that code for the N-terminal LIM domain known to interact with proteins that link the actin cytoskeleton to the cell membrane(Zhang et al. 2001). The LIM domain is also responsible for *Nrap*’s role in myofibrillogenesis during development(Carroll et al. 2001). Alternative splicing of exon 2 may modulate the binding affinity of the *Nrap* LIM domain to these myofibrillar and linker proteins.

At 22 kb, nebulin is over twice the length of IsoSeq’s maximum read length. This resulted in reads of partial nebulin transcripts. However, internal priming enabled tiled coverage across the entire length of the transcripts. We only detected alternative splicing of the 3’ half of the gene encoding the super repeat and Z-disk regions of the protein. As these alternatively spliced exons are close enough in proximity, they fit within a single read and were phased together. In silico translation of mouse nebulin used an older and more inclusive annotation containing 166 exons which is most similar to ENSMUST00000238749.1 in Gencode release M22 with 165 exons(Buck et al. 2010). However, this transcript is likely not accurate as it includes both exons 127 and 128, which are expressed in a mutually exclusive manner(Donner et al. 2006; Lam et al. 2018). Thus, we used the only mouse nebulin RefSeq annotation (NM_010889.1/ENSMUST00000036934.11) that contains 157 exons as a reference and below we clarify the annotation further.

Using our exon-based analysis on IsoSeq data, we detected differential splicing of multiple nebulin exons between 122-152 (NM_010889.1). Exons 13-132 (13-137 of ENSMUST00000238749.1) code for the super repeat region and exons 134-157 (139-165) code for the Z-disk region of nebulin in mouse. The majority of differential splicing of nebulin was detected within the Z-disk region which included an unannotated exon u-002 (Figure 1, 4A, and 6C). Exon u-002 is in-frame and conserved in human nebulin transcripts (see footnote). In the fast-twitch EDL, most of the exons of the Z-disk region are removed while the slow-twitch soleus included the majority of exons at a higher percentage.

The Z-disk plays a key role in contractile force transmission within and across sarcomeres. Differential splicing of the nebulin Z-disk region correlates with Z-disk width being thinner in white muscles and thicker in red muscles(Rowe 1973). Independent observations also found that the slow soleus and fast tibialis cranialis muscles corroborate our splicing data, suggesting that nebulin may have a functional role regulating Z-disk width(Buck et al. 2010).

We also detected alternative splicing of exon 127 and 128 (ENSMUST00000238749.1), which are located in the super repeat region right outside of the Z-disk anchorage point. They are expressed in a mutually exclusive manner, are conserved in humans (NM_001271208), and code for peptides with different charge and hydrophobicity, suggesting distinctive functional roles(Donner et al. 2006; Lam et al. 2018). In humans, these exons show developmentally regulated inclusion(Lam et al. 2018). The peptides in the super repeat region of nebulin are binding sites for kelch-like family member 40 (*KLHL40*), loss of which causes a nemaline-like myopathy(Garg et al. 2014). In turn, *KLHL40* is suggested to stabilize nebulin and Leimodin 3 (*LMOD3*, another protein implicated in nemaline myopathy) by acting like a chaperone and maintaining proper folding of these two proteins during muscle contraction(Garg et al. 2014).

Using exPhaser we identified and quantified multiple novel nebulin transcripts in both the Z-disk and 3’ end of super repeat region (Figure 6C). Thus, our phasing data presents a catalog of more rigorous splice combinations. It is worth noting that none of the current nebulin transcripts annotated in RefSeq and Gencode release M10 (see footnote), are expressed in the EDL and Soleus muscles.

Titin transcripts (∼106 kb) are over tenfold longer than maximum IsoSeq read lengths. This did not thwart the ability to determine differential exon usage through our exon-based exCOVator analysis. However, the maximum read length put limitations on the exPhaser analysis. The phasing analysis required breaking up the exons into multiple groups based on proximity. We then quantified transcripts known to be cardiac and skeletal muscle specific in mouse. This helped uncover multiple novel splicing patterns expressed in specific muscle types. Using this strategy together with Cardiodb, TITINdb and other resources(Savarese et al. 2018; Roberts et al. 2015; Laddach et al. 2017), we compared our mouse findings to human studies to determine if alternative splicing of these exons are conserved.

Cardiac specific cassette exon 191 (NM_133378, N2-A) in human (243 LRG) codes for one of multiple immunoglobulin-like (Ig-like) domains in the highly extensible I-band region of titin(Linke 2017). Ig-like domains can quickly unfold and refold, a property important for muscle elasticity and passive stiffness(Rief et al. 1997). This is especially relevant in heart where stiffness can affect the ejection fraction. It’s possible that this cassette exon is differentially spliced to adjust the length of the I-band region of titin and stiffness of sarcomeres in the heart(Opitz et al. 2004).

Exon 312 in mouse or 363 (LRG) in human is the penultimate exon of titin. This c-terminal region integrates into the M-band of the sarcomere, functions as a scaffold and proposed to play a role in mechanosensitivity(Linke 2017; Zacharchenko et al. 2015). Exon 363 (LRG) codes for the N-terminal portion of the final Ig domain M10 (TitinDb). Mutations in M10 cause two types of muscular dystrophies (OMIM) as it is important for interacting with obscurin and myospryn in the M-line(Sarparanta et al. 2010). Specific alternative splicing of exon 363 (LRG) in skeletal muscle may alter titin’s protein interactions in the M-line.

Alternative 3’ exon 45 (exon 48 LRG) is part of the smallest titin isoform conserved in humans as Novex-3. The isoform is known to interact with obscurin in the I-band and a variety of proteins in the Z-disk(Bang et al. 2001). It is speculated to play a role in myofibrillar signaling during muscle development and cardiac disease(Bang et al. 2001). Novex-3 is expressed in all striated muscles, but not smooth muscles(Bang et al. 2001). It is interesting that we found it expressed more in skeletal muscle compared to cardiac muscle as it is often labeled as a cardiac isoform.

Exons 11, 12 and 13 of titin are conserved in humans with the same numbering (LRG). They code for z-repeat domains 4, 5, and 6 respectively in the N-terminal region of titin imbedded in the Z-disk. These exons are variably spliced in human and may be involved in assembling Z-disks of variable width(Gautel et al. 1996). Titin’s Z-repeats interact with alpha-actinin in the Z-disk, and have been proposed to be important for Z-disk organization, assembly and force transmission between adjacent sarcomeres(Joseph et al. 2001). Alternative splicing of these exons seems to correlate with Z-disk width much like the Z-disk region of nebulin; transcripts produced by the fast EDL muscle excludes more exons compared to Soleus. In addition, exon 11 of titin is also always spliced out in skeletal muscle in a similar manner to humans(Savarese et al. 2018).

IsoSeq is generally recommended for transcript isoform discovery rather than for quantitative measures. In contrast, we found that counting full-length reads extracted from uncollapsed cluster reads was quantitative based on our validations. However, there are false positives caused by spurious reads that require manual inspection in a genome browser.

Recent improvements to sample preparation and data processing in IsoSeq 3 (SmrtLink v8) allow improved read length and data quality. The use randomized primers could also open the door for more evenly distributed coverage for ultra-long transcripts but would require some changes in the data processing pipeline.

Our study analyzed alternative splicing at a bulk tissue level, but it is unknown if the alternatively spliced transcripts are expressed by different muscle fibers in the muscle or by the same fiber. Use of single-cell isoform sequencing approaches that combine single-cell RNA-seq with IsoSeq would be needed to answer these questions(Gupta et al. 2018). Further, new methods for isolating intact ultra-long transcripts and converting them to cDNA would improve alternative splicing analysis of large structural genes. Improved direct RNA sequencing methods being developed may also help circumvent this limitation(Garalde et al. 2018).

We successfully applied our strategy to study three important muscle structural genes that produce complex, repetitive, ultra-long transcripts. It is difficult at best, to assess the alternative splicing of these transcripts with short-read data, and current long-read analysis approaches, are also not yet optimized. Our strategy provides a potential solution to these issues and enables simultaneous ultra-long transcript isoform discovery and quantitation, opening the door to differential expression analysis of difficult transcripts.

## FOOTNOTE

Our analysis was based on Gencode release M10 (Ensembl 85) which matches the latest Gencode genome reference used to map the IsoSeq data. At the time of writing, we checked the latest genode annotation (release M22, Ensembl 97) to make sure that our findings were up to date. We found no annotation changes that would affect our analysis of titin and *Nrap*.

However, we found that two new inferred transcripts (ENSMUST00000238749.1 and ENSMUST00000238288.1) were added to nebulin in Gencode release M22 that contain all exons found to be missing in the RefSeq annotation NM_010889.1 (including the unannotated u-002). The three exons between 147 and 148 labeled as 3*, u-002 and 143** in figure 6C are annotated in M22 as 153, 154, and 155 of ENSMUST00000238749.1 respectively. The two mutually exclusive exons labeled as 7*** and 123 in figure 6C are 127 and 128 of ENSMUST00000238749.1 respectively. The four exons found between 85 and 86 in NM_010889.1 are annotated in release M22 as exons 71,72,73 and 74 of ENSMUST00000238749.1. In the phasing results (figure 6C), the nebulin “UNANNOTATED 7” transcript matches the current annotation ENSMUST00000238288.1 (exon 128 spliced out).

Interestingly, this is still not the highest expressed transcript in the Soleus (UNANNOTATED 8 at 53.2% of 817 FL-reads). The annotation ENSMUST00000238749.1 (containing all exons) is not expressed in any of the muscles further supporting that exons 127 and 128 are mutually exclusive and the annotation is likely incorrect but would be useful as a meta or inferred complete transcript similar to NM_001267550.1 (ENST00000589042).

## METHODS

### Animals and tissue harvesting

Six-week old C57BL/6 (WT) male mice (JaxLabs) were euthanized using CO2 followed by cervical dislocation. EDL, Soleus and cardiac muscles from the same animal were excised. For cardiac muscle, only 10 mg of the apex was acquired. The excised muscles were immediately flash frozen in 2-methybutane (Sigma) in liquid nitrogen and stored at −80°C for further processing.

### RNA isolation and purification

The cardiac apex was processed individually, while the left and right EDL, and Soleus from the same mouse were pooled by muscle type for processing. The muscles were thawed to −20°C, cut into 2 mm cubes, placed into pre-chilled T-prep devices (COVARIS, Cat# 520097) and crushed into a fine powder using the impactor (COVARIS, Cat# 500305) following manufacturer’s protocol. Afterwards, 600μl of lysis buffer from the MirVana RNA Isolation kit (Thermo Fisher: AM1560) was added to each T-prep device, the homogenate transferred into fresh tubes and total RNA was extracted following the MirVana protocol.

DNA was removed using the Turbo DNA-free kit (ThermoFisher, #AM1907). Purification by ethanol precipitation was performed by mixing 0.1x volume of 3M sodium acetate (ThermoFisher, #AM9740) and 3x volume of 100% molecular grade ethanol into the RNA samples. The mixture incubated overnight at −20°C and centrifuged at 10,000g at 4°C for 20 minutes. The supernatant discarded, and RNA pellet rinsed with 400μl of fresh 70% ethanol and centrifuged again. This wash step was repeated twice before the RNA pellet was left to dry at room temperature for 5 minutes. The pellets were resuspended in nuclease-free water and assessed for purity using a NanoDrop 2000. All samples had 260/280 ratios > 1.8 and 260/230 > 2.0

Assessing RNA quantity and quality. RNA was quantified using a Qubit Fluorometer (v2.0) and RNA broad range assay kit (ThermoFisher, #Q10210). Quality of total RNA was assessed on an Agilent BioAnalyzer 2100 using the RNA 6000 Nano kit (Agilent, # 5067-1511). All samples had a RIN > 7.6. The purified total RNA samples were stored in −80°C.

### IsoSeq library generation and sequencing

Total RNAs from one mouse was reverse transcribed following PacBio’s IsoSeq procedure and checklist using Clonetech SMARTer PCR cDNA Synthesis Kit and BluePippin Size-Selection System (P/N100-377-100-04). Three starting reactions of 600ng-1μg total RNA were used per muscle sample for first stand cDNA synthesis. Custom 3’ primers containing barcodes were added during this step: soleus (dT_BC1), EDL (dT_BC2), and Cardiac (dT_BC3) following the IsoSeq Barcoding protocol. After first strand synthesis, the number of cycles chosen for large scale PCR were: 12 for soleus and EDL and 10 for cardiac samples.

After amplification, cDNA libraries were pooled in equimolar amounts and the subsequent steps were performed according to protocol with modifications. During the BluePippin size selection step, 500ng of cDNA was loaded per lane of the gel cassette and only the 5-10kb size fraction were collected for further library generation. SMRTbell adapters were added using the SMRTbell Template Prep Kit 1.0 (#100-259-100). A second BluePippin size selection step was performed after SMRTbell templates were generated to remove small library fragments.

Single-molecule sequencing. Sequencing was performed according to PacBio RS II protocol. Briefly, DNA polymerase was bound using DNA/Polymerase Binding Kit P6 V2 (#100-372-700). DNA P6 Control Complex (#100-356-500) was added and DNA Sequencing Reagent 4.0 (#100-356-200) was loaded onto the system. During sequencing loading optimization (first 8 SMRTcells), two concentrations were used on four cells each: 1.0nM and 1.2nM. The 1.2nM concentration resulted in a higher yield of total reads of interest and the concentration was used for the remaining SMRTcells. The pooled IsoSeq libraries were sequenced using a total of 24 SmrtCells (#100-171-800) on the PacificBiosciences RSII machine (output statistics in Table S1).

### Data processing and alignment

Raw data (bax.h5) was processed according to the official Iso-Seq protocol using SmrtAnalysis (v2.3) with a few modifications to help process the large amount of data. Briefly, the fasta files produced from individual SmrtCells after the circular consensus step (CCS) were merged into two fasta files composed of eight SmrtCells. The two merged files were demultiplexed into the three muscle samples based on their barcodes yielding six total files. Barcodes numbers were converted to zero-based according to protocol (dT_BC1àBC0, BC1àdT_BC2, and dT_BC3àBC2). The six files were processed through classify and cluster steps in the IsoSeq method to produce fastq files. Fastq files belonging to each muscle were concatenated producing three large fastq files (e.g. soleus.fastq, edl.fastq, cardiac.fastq) ready for alignment. To allow for this type of merging, cluster numbers (c1, c2, c3 etc) from each fastq file’s read names were renumbered for each consecutive fastq file to avoid overwriting redundant cluster numbers. See mergeFastq.py script in github repository. Finally, the alignment steps were performed using STAR v2.5.0a and GENCODE primary mouse sequence and annotation (GRCm38) version M10 (Ensembl 85) released 2016-07-19. All data processing steps were performed on the Colonial One George Washington University HPC using the Slurm workload manager.

### Reverse-Transcription PCR and Sanger Sequencing

Primers were designed using NCBI Primer-Blast (Table S2). RT-PCR reactions were performed using SuperScript III One-Step RT-PCR system (ThermoFisher, #12574030) following manufacturing protocol. Samples used in validation (mouse 2 and 3) are independent from the sequenced sample (mouse1). The amplification products were loaded into a 2% agarose gel made with 1x TBE buffer and 5μl ethidium bromide for electrophoresis and imaging.

The RT-PCR products were excised from the agarose gel and purified using Qiagen gel purification kit and sent along with primers to Quintarabio for Sanger Sequencing. Alignments of the sequences were visualized in IGV after using the BLAT tool.

### Bioinformatics analysis

Scripts for identifying novel exons, differential exon usage analysis, exon phasing and transcript structure quantification were created using python and the HT-Seq library. For full details of the analysis approach see supplementary methods and the GitHub page (see data availability section).

### DATA ACCESS

All raw and processed sequencing data generated in this study have been submitted to the NCBI Gene Expression Omnibus (GEO; https://www.ncbi.nlm.nih.gov/geo/) under accession number GSE138362. All scripts, detailed notes, links and archived versions of PacBio protocols used in this project are made available in our GitHub repository (https://github.com/puapinyoying/isoseq_manuscript_resources).

## Supporting information

exPhaser_implementation

## ACKNOWLEDGEMENT

We would like to thank Adam K. L. Wong, Keith Crandall and the Colonial One HPC team at George Washington university for providing technical support and computational resources for the project. Hiroki Morizono for bioinformatics lessons and support. Linda Werling for writing and student support. Healther Locovare for helping us optimize the IsoSeq bench protocol for our project. We would also like to thank past and present members of the Eric Hoffman lab and Genetic Medicine department at Children’s National Hospital for helpful discussions. This work was supported by the National Institutes of Health R01NS029525.

## REFERENCES

1. Bang M-L, Centner T, Fornoff F, Geach AJ, Gotthardt M, McNabb M, Witt CC, Labeit D, Gregorio CC, Granzier H, et al. 2001. The Complete Gene Sequence of Titin, Expression of an Unusual 700-kDa Titin Isoform, and Its Interaction With Obscurin Identify a Novel Z-Line to I-Band Linking System. Circ Res 89: 1065–1072.

2. Bang M, Chen J. 2015. Roles of Nebulin Family Members in the Heart. Circ J 79: 2081–7.

3. Buck D, Hudson BD, Ottenheijm CAC, Labeit S, Granzier H. 2010. Differential splicing of the large sarcomeric protein nebulin during skeletal muscle development. J Struct Biol 170: 325–333.

4. Buck D, Weirather JL, de Cesare M, Wang Y, Piazza P, Sebastiano V, Wang XJ, Au KF. 2017. Comprehensive comparison of Pacific Biosciences and Oxford Nanopore Technologies and their applications to transcriptome analysis. F1000Research 6.

5. Carroll S, Lu S, Herrera AH, Horowits R. 2004. N-RAP scaffolds I-Z-I assembly during myofibrillogenesis in cultured chick cardiomyocytes. J Cell Sci 117: 105–14.

6. Carroll SL, Herrera a H, Horowits R. 2001. Targeting and functional role of N-RAP, a nebulin-related LIM protein, during myofibril assembly in cultured chick cardiomyocytes. J Cell Sci 114: 4229–38.

7. Carroll SL, Horowits R. 2000. Myofibrillogenesis and formation of cell contacts mediate the localization of N-RAP in cultured chick cardiomyocytes. Cell Motil Cytoskeleton 47: 63–76.

8. Castle JC, Zhang C, Shah JK, Kulkarni A V., Kalsotra A, Cooper TA, Johnson JM. 2008. Expression of 24,426 human alternative splicing events and predicted cis regulation in 48 tissues and cell lines. Nat Genet 40: 1416–1425.

9. Cummings BB, Marshall JL, Tukiainen T, Lek M, Donkervoort S, Foley AR, Bolduc V, Waddell L, Sandaradura S, O’Grady GL, et al. 2017. Improving genetic diagnosis in Mendelian disease with transcriptome sequencing. Sci Transl Med 5209: 074153.

10. Dhume A, Lu S, Horowits R. 2006. Targeted disruption of N-RAP gene function by RNA interference: A role for N-RAP in myofibril organization. Cell Motil Cytoskeleton 63: 493– 511.

11. Donner K, Nowak KJ, Aro M, Pelin K, Wallgren-Pettersson C. 2006. Developmental and muscle-type-specific expression of mouse nebulin exons 127 and 128. Genomics 88: 489–495.

12. Donner K, Sandbacka M, Lehtokari V-L, Wallgren-Pettersson C, Pelin K. 2004. Complete genomic structure of the human nebulin gene and identification of alternatively spliced transcripts. Eur J Hum Genet 12: 744–51.

13. Garalde DR, Snell EA, Jachimowicz D, Sipos B, Lloyd JH, Bruce M, Pantic N, Admassu T, James P, Warland A, et al. 2018. Highly parallel direct RNA sequencing on an array of nanopores. Nat Methods 15: 201–206.

14. Garg A, O’Rourke J, Long C, Doering J, Ravenscroft G, Bezprozvannaya S, Nelson BR, Beetz N, Li L, Chen S, et al. 2014. KLHL40 deficiency destabilizes thin filament proteins and promotes nemaline myopathy. J Clin Invest 124: 3529–39.

15. Gautel M, Goulding D, Bullard B, Weber K, Fürst DO. 1996. The central Z-disk region of titin is assembled from a novel repeat in variable copy numbers. J Cell Sci 109 ( Pt 1: 2747–54.

16. Guo W, Bharmal SJ, Esbona K, Greaser ML. 2010. Titin diversity--alternative splicing gone wild. J Biomed Biotechnol 2010: 753675.

17. Gupta I, Collier PG, Haase B, Mahfouz A, Joglekar A, Floyd T, Koopmans F, Barres B, Smit AB, Sloan SA, et al. 2018. Single-cell isoform RNA sequencing characterizes isoforms in thousands of cerebellar cells. Nat Biotechnol 36.

18. Herrera AH, Elzey B, Law DJ, Horowits R. 2000. Terminal regions of mouse nebulin: Sequence analysis and complementary localization with N-RAP. Cell Motil Cytoskeleton 45: 211.

19. Hoffman EP, Kunkel LM. 1989. Dystrophin abnormalities in Duchenne/Becker muscular dystrophy. Neuron 2: 1019–29.

20. Joseph C, Stier G, O’Brien R, Politou AS, Atkinson RA, Bianco A, Ladbury JE, Martin SR, Pastore A. 2001. A structural characterization of the interactions between titin Z-repeats and the α-actinin C-terminal domain. Biochemistry 40: 4957–4965.

21. Labeit S, Gibson T, Lakey A, Leonard K, Zeviani M, Knight P, Wardale J, Trinick J. 1991. Evidence that nebulin is a protein-ruler in muscle thin filaments. FEBS Lett 282: 313–316.

22. Labeit S, Kolmerer B. 1995. The complete primary structure of human nebulin and its correlation to muscle structure. J Mol Biol 248: 308–315.

23. Laddach A, Gautel M, Fraternali F. 2017. TITINdb-a computational tool to assess titin’s role as a disease gene. Bioinformatics 33: 3482–3485.

24. Lam LT, Holt I, Laitila J, Hanif M, Pelin K, Wallgren-Pettersson C, Sewry CA, Morris GE. 2018. Two alternatively-spliced human nebulin isoforms with either exon 143 or exon 144 and their developmental regulation. Sci Rep 8: 15728.

25. Li Y, Rao X, Mattox WW, Amos CI, Liu B. 2015. RNA-seq analysis of differential splice junction usage and intron retentions by DEXSeq. PLoS One 10: 1–14.

26. Linke WA. 2017. Titin Gene and Protein Functions in Passive and Active Muscle. Annu Rev Physiol 80: 389–411.

27. Lu S, Borst DE, Horowits R. 2008. Expression and alternative splicing of N-RAP during mouse skeletal muscle development. Cell Motil Cytoskeleton 65: 945–54.

28. Luo G, Zhang JQ, Nguyen TP, Herrera AH, Paterson B, Horowits R. 1997. Complete cDNA sequence and tissue localization of N-RAP, a novel nebulin-related protein of striated muscle. Cell Motil Cytoskeleton 38: 75–90.

29. McElhinny AS, Schwach C, Valichnac M, Mount-Patrick S, Gregorio CC. 2005. Nebulin regulates the assembly and lengths of the thin filaments in striated muscle. J Cell Biol 170: 947–57.

30. Mohiddin SA, Lu S, Cardoso JP, Carroll S, Jha S, Horowits R, Fananapazir L. 2003. Genomic organization, alternative splicing, and expression of human and mouse N-RAP, a nebulin-related LIM protein of striated muscle. Cell Motil Cytoskeleton 55: 200–212.

31. Nadal-Ginard B, Smith CW, Patton JG, Breitbart RE. 1991. Alternative splicing is an efficient mechanism for the generation of protein diversity: contractile protein genes as a model system. Adv Enzyme Regul 31: 261–86.

32. Nakka K, Ghigna C, Gabellini D, Dilworth FJ. 2018. Diversification of the muscle proteome through alternative splicing. 1–18.

33. Nam DK, Lee S, Zhou G, Cao X, Wang C, Clark T, Chen J, Rowley JD, Wang SM. 2002. Oligo(dT) primer generates a high frequency of truncated cDNAs through internal poly(A) priming during reverse transcription. Proc Natl Acad Sci U S A 99: 6152–6.

34. Nilsen TW, Graveley BR. 2010. Expansion of the eukaryotic proteome by alternative splicing. Nature 463: 457–463.

35. Opitz CA, Leake MC, Makarenko I, Benes V, Linke WA. 2004. Developmentally regulated switching of titin size alters myofibrillar stiffness in the perinatal heart. Circ Res 94: 967–75.

36. Pappas CT, Bliss KT, Zieseniss A, Gregorio CC. 2011. The Nebulin family: an actin support group. Trends Cell Biol 21: 29–37.

37. Pistoni M, Ghigna C, Gabellini D. 2013. Alternative splicing and muscular dystrophy. 7: 441– 452.

38. Politou AS, Spadaccini R, Joseph C, Brannetti B, Guerrini R, Helmer-Citterich M, Salvadori S, Temussi PA, Pastore A. 2002. The SH3 domain of nebulin binds selectively to type II peptides: theoretical prediction and experimental validation. J Mol Biol 316: 305–15.

39. Rief M, Gautel M, Oesterhelt F, Fernandez JM, Gaub HE. 1997. Reversible unfolding of individual titin immunoglobulin domains by AFM. Science 276: 1109–12.

40. Roberts AM, O’Regan DP, De Marvao A, Dawes TJW, Cook SA, Ware JS, Baksi J, Buchan RJ, Walsh R, John S, et al. 2015. Integrated allelic, transcriptional, and phenomic dissection of the cardiac effects of titin truncations in health and disease. Sci Transl Med 7.

41. Rowe RW. 1973. The ultrastructure of Z disks from white, intermediate, and red fibers of mammalian striated muscles. J Cell Biol 57: 261–77.

42. Sarparanta J, Blandin G, Charton K, Vihola A, Marchand S, Milic A, Hackman P, Ehler E, Richard I, Udd B. 2010. Interactions with M-band titin and calpain 3 link myospryn (CMYA5) to tibial and limb-girdle muscular dystrophies. J Biol Chem 285: 30304–30315.

43. Savarese M, Jonson PH, Huovinen S, Paulin L, Auvinen P, Udd B, Hackman P. 2018. The complexity of titin splicing pattern in human adult skeletal muscles. Skelet Muscle 8: 1–9.

44. Sebastian S, Faralli H, Yao Z, Rakopoulos P, Palii C, Cao Y, Singh K, Liu QC, Chu A, Aziz A, et al. 2013. Tissue-specific splicing of a ubiquitously expressed transcription factor is essential for muscle differentiation. Genes Dev 27: 1247–1259.

45. Steijger T, Abril JF, Engström PG, Kokocinski F, RGASP Consortium, Hubbard TJ, Guigó R, Harrow J, Bertone P. 2013. Assessment of transcript reconstruction methods for RNA-seq. Nat Methods 10: 1177–84.

46. Trapnell C, Williams Ba, Pertea G, Mortazavi A, Kwan G, van Baren MJ, Salzberg SL, Wold BJ, Pachter L. 2010. Transcript assembly and quantification by RNA-Seq reveals unannotated transcripts and isoform switching during cell differentiation. Nat Biotechnol 28: 511–5.

47. Treangen TJ, Salzberg SL. 2011. Repetitive DNA and next-generation sequencing: computational challenges and solutions. Nat Rev Genet 13: 36–46.

48. Wang B, Tseng E, Regulski M, Clark TA, Hon T, Jiao Y, Lu Z, Olson A, Stein JC, Ware D. 2016. Unveiling the complexity of the maize transcriptome by single-molecule long-read sequencing. Nat Commun 7: 11708.

49. Wang ET, Sandberg R, Luo S, Khrebtukova I, Zhang L, Mayr C, Kingsmore SF, Schroth GP, Burge CB. 2008. Alternative isoform regulation in human tissue transcriptomes. Nature 456: 470–6.

50. Wang K, Wright J. 1988. Architecture of the sarcomere matrix of skeletal muscle: immunoelectron microscopic evidence that suggests a set of parallel inextensible nebulin filaments anchored at the Z line. J Cell Biol 107: 2199–212.

51. Wang Z, Gerstein M, Snyder M. 2009. RNA-Seq: a revolutionary tool for transcriptomics. Nat Rev Genet 10: 57–63.

52. Witt CC, Burkart C, Labeit D, McNabb M, Wu Y, Granzier H, Labeit S. 2006. Nebulin regulates thin filament length, contractility, and Z-disk structure in vivo. EMBO J 25: 3843–3855.

53. Yang X, Coulombe-Huntington J, Kang S, Sheynkman GM, Hao T, Richardson A, Sun S, Yang F, Shen YA, Murray RR, et al. 2016. Widespread Expansion of Protein Interaction Capabilities by Alternative Splicing. Cell 164: 805–17.

54. Zacharchenko T, von Castelmur E, Rigden DJ, Mayans O. 2015. Structural advances on titin: towards an atomic understanding of multi-domain functions in myofilament mechanics and scaffolding. Biochem Soc Trans 43: 850–855.

55. Zhang JQ, Elzey B, Williams G, Lu S, Law DJ, Horowits R. 2001. Ultrastructural and biochemical localization of N-RAP at the interface between myofibrils and intercalated disks in the mouse heart. Biochemistry 40: 14898–14906.

