## Supplementary figures and methods for "A new long-read RNA-seq analysis approach identifies and quantifies novel transcripts of very large genes"

### SUPPLEMENTARY DATA INDEX

- Supplementary figures and tables
- Supplementary methods
- exPhaser implementation (separate pdf)

### SUPPLEMENTARY FIGURES AND TABLES

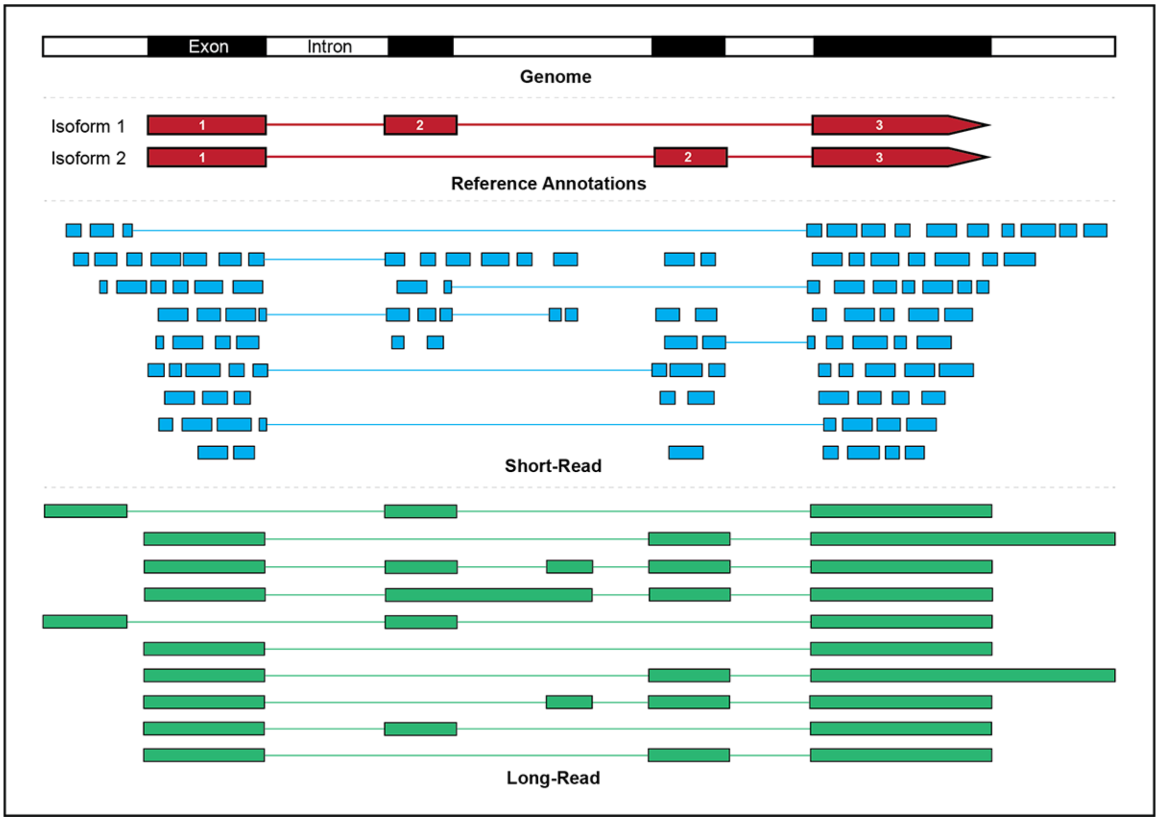


Supplementary Figure 1. Comparison between short-read and long-read RNA-Sequencing. Short-reads (<=300bp) are too short to span more than two exon junctions. Algorithms are used to assemble transcripts from the data. However, predictions based on data can miss or incorrectly predict novel isoforms from mis-mapped reads and artifacts. Long-read sequencing (<= 10kb+, average of 6kb) may span entire transcripts or large portions of them which allows for phasing of exons and isoform identification.

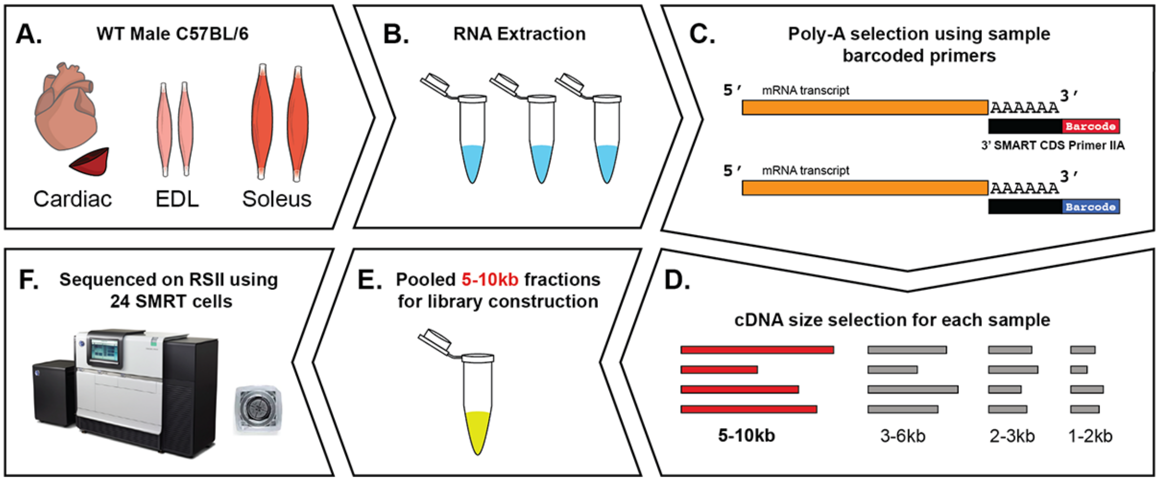


Supplementary Figure 2. Experimental design and sequencing approach. A) Cardiac apex, extensor digitorum longus (left and right), and soleus (left and right) muscles were surgically removed from wild-type male C57BL/6 mice. B) Total RNA was extracted from each muscle. C) cDNAs from each muscle sample were generated using barcoded Oligo-dT primers (3' SMRT CDS Primer IIA). D) Next cDNAs were size selected for 5-10kb fraction using the BluePippen system. E) the 5-10kb fractions from all three muscles were pooled for library construction. F) Library was sequenced on a PacBio RSII system using 24 SMRT Cells.

**Supplementary Table 1. Sequencing statistics from classify output (24 SMRT cells)**

| Multiplexed (pooled) | Number of reads |
|---|---|
| Reads of Insert (ROI) | 1,955,502 |
| Five prime (5') | 711,096 |
| Three prime (3') | 1,020,887 |
| Poly-A | 935,179 |
| Filtered short reads | 26,521 |
| Non-full-length | 1,415,922 |
| Full-length* | 513,059 |
| Full-length non-chimeric** | 509,095 |

| Demultiplexed | Reads of insert (ROI) |
|---|---|
| Soleus (barcode 0) | 163,368 |
| EDL (barcode 1) | 167,632 |
| Cardiac (barcode 2) | 178,095 |
| Total identified by barcode | 509,095 |

| Average read length | Base pairs |
|---|---|
| Full-length non-chimeric | 6,009 |

***Full-length** - determined by presence of cDNA library primers and poly-A sequence
****Chimeric reads** (artificial chimeras) - sequencing artifacts formed by insufficient SMRTbell adapters, random fusion of ligated transcripts during PCR or true biological fusion events of two separate transcripts that cannot be distiguished

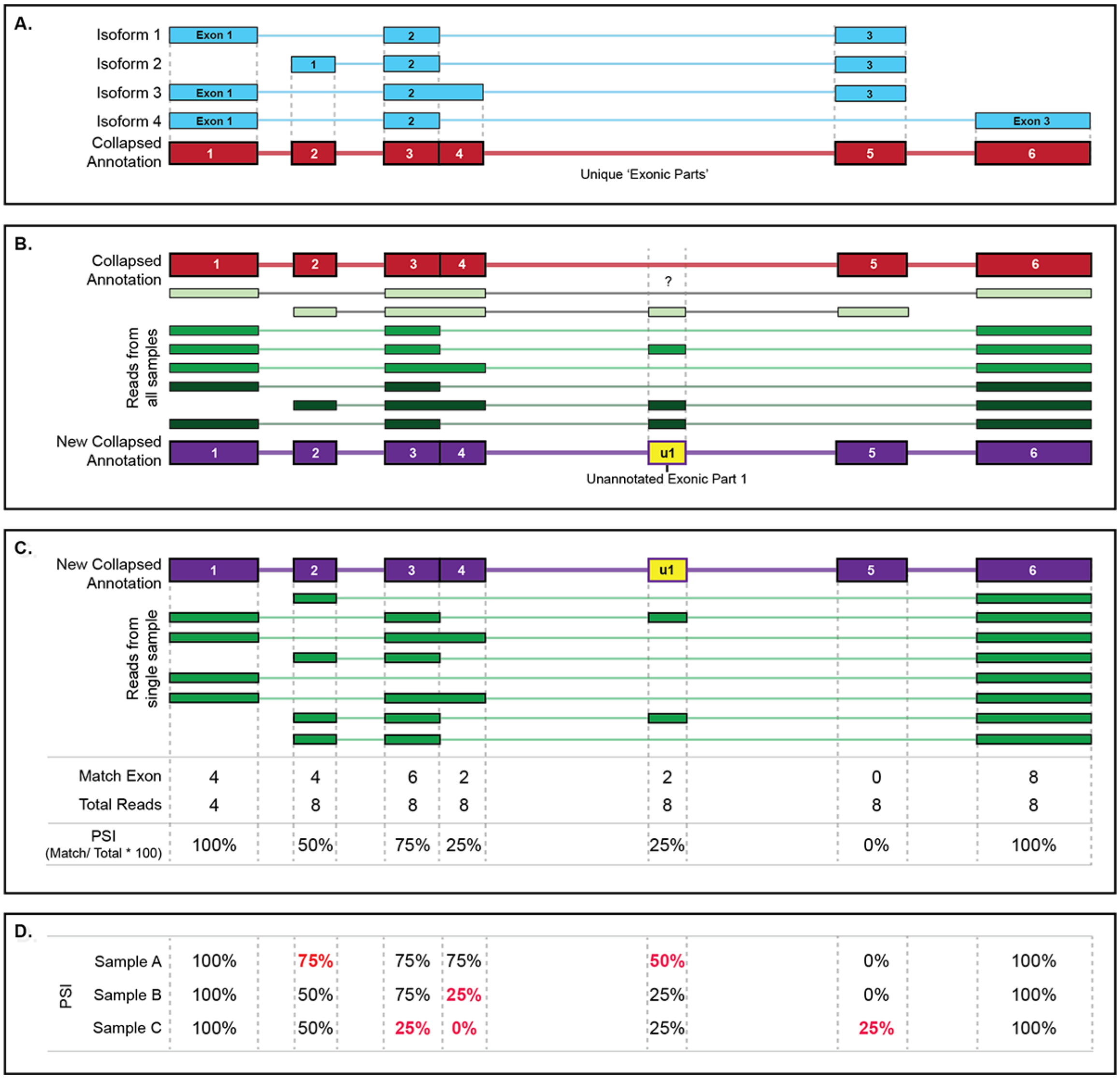


Supplementary Figure 3. Custom exon-based IsoSeq analysis pipeline (exCOVater). A) Transcript isoforms often share multiple exons complicating analysis. Using the 'dexseq_prepare_annotation.py' script, annotations were collapsed into unique exonic parts (EP) for analysis. B) Read data from all samples are used to find unannotated sequences (potential novel exons) not present in the collapsed annotation file. Unannotated exons (e.g. u1) are added to the list of annotations for differential usage analysis. C) Count match and total reads for each all EPs for individual samples and calculate a ratio (match/total) that is multiplied by 100 to obtain the percent spliced-in (PSI). In this illustration, EP6 is constitutively expressed; EP1 and EP2 are alternate 5’ exons; EP5 is an alternate 3’ exon that is not expressed in this sample; EP3, EP4 and unannotated exonic part 1 (u1) are cassette exons. D) Observing differential exon usage across samples. The ratio is a normalized value that can be used to compare exon usage across multiple samples. The ratios for each sample are used by ‘filterDiffUsedExons.py’ for further filtering exons with less than defined % difference across samples. EP1 and EP6 have 0% change across samples and would be filtered out. EP2-5 and u1 are differentially used and will be retained for further analysis.

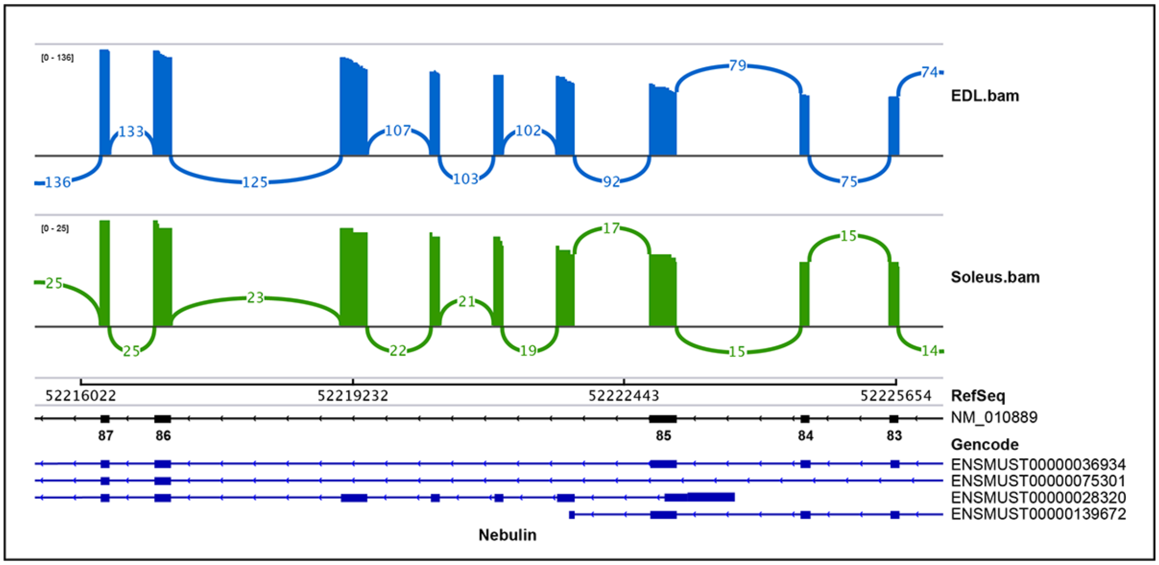


Supplementary Figure 4. Sashimi plot showing the splicing pattern of Nebulin exons 83-87 in EDL and Soleus muscles. The data shows 4 exons between 85 and 86 that are not annotated in the RefSeq database and are constitutively expressed in both EDL and Soleus skeletal muscles. These 4 unannotated exons are represented in Gencode M10 (seen in figure) in ENSMUST00000028320 and Gencode M22 in two additional transcripts ENSMUST00000238749 and ENSMUST00000238288 (not shown).

**Supplementary Table 2. Primers used for RT-PCR and Sanger Sequencing validation**

| Gene | Exon | Exon Location | RefSeq (mm10) | Primer Dir | Primed Exons | Primer sequence 5'-->3' |
|---|---|---|---|---|---|---|
| *Neb* | 138 | chr2:52165167-52165277 | NM_010889.1 | Fwd | 134-135 | ACCTTGCAAGCGAGGTCAA |
| *Neb* | 138 | chr2:52165167-52165277 | NM_010889.1 | Rev | 139 | GCTCTCAGCATGTCAGGAGT |
| *Nrap* | 12 | chr19:56374364-56374468 | NM_008733.4 | Fwd | 9 | GACCGATGTGGCCAGGTTTACTCAGAAG |
| *Nrap* | 12 | chr19:56374364-56374468 | NM_008733.4 | Rev | 14 | CAGGGGAACCAGCCTCATCGTTGTTTG |
| *Ttn* | 191 | chr2:76802231-76802497 | NM_011652.3 | Fwd | 189-190 | TGGAGCCTAACGATAAGGTGG |
| *Ttn* | 191 | chr2:76802231-76802497 | NM_011652.3 | Rev | 194 | CATCCTCAAGCTGCGCATTC |

### SUPPLEMENTARY METHODS

#### exCOVator exon-based analysis pipeline

*Identifying novel exons and differential usage analysis.* The goal of this analysis pipeline was to compare splice differences between the three different tissues and identify unannotated / novel exons. We decided to compare differential exon usage (inspired by DEXSeq for short reads) between samples. This approach can measure relative exon usage across conditions regardless if the full-length reads cover entire transcripts or if the read was derived from internal priming. Therefore, analysis through differential exon usage would open up opportunities to analyze and compare the splicing behavior of extremely long transcripts such as ones from structural genes that are far beyond technical sequencing limitations.

*Counting number of cluster-reads per gene***.** During the preliminary analysis, determining which muscle structural genes to observe consisted of randomly choosing familiar gene names into the IGV browser. We need a systematic way to narrow down the scope of the analysis. According to the Gencode release archive for version M10, there are 48,440 total functional elements in the genome. These functional elements are discrete units or genes located on the genome. We needed to determine which of these genes were transcribed and were sequenced. In other words, how many of these genes contain sequence coverage or cluster reads mapped to them. In order to determine which genes had mapped reads, the number of cluster-reads per gene in each alignment (BAM) file were counted using HTSeq-count software. We found 3219 genes that had at least one cluster read in one of the three samples. This step reduced the scope of the analysis to 6.65% of the total genes annotated in the Gencode file. Since we sequenced a specific tissue type (muscle), not all of the genes in the genome should be expressed (contain no RNA transcripts or mapped reads). By reducing the number of target genes to a fraction of the total, we were able to dramatically reduce noise and computational load for downstream analyses.

*Obtaining a list of genes with coverage above a minimum threshold.* he next step was to generate a list of genes that had enough cluster reads to be confident in the data and be able to compare differential exon usage between samples. To do this, the cluster read count data was imported into Excel. We chose a minimum cut-off filter of 10 reads per gene in at least one of the three muscle samples. This method yielded 1520 genes that met the minimum threshold minus the Riken genes from the data set (see 'excovator_exon-based_counts.xlsx' in the processed data found in the GEO repository for this manuscript). The final result was a list of 686 unique genes with >= 10 cluster reads present in at least one of the three samples. These criteria were selected because some genes may be uniquely expressed in one of the muscle types or poorly expressed in some tissues (e.g. Nebulin in heart). This method ensured that those genes would not be filtered out of the analysis.

*Reducing the number of annotations to analyze from the (Gencode) GTF file.* Once We obtained the list of 686 genes to analyze, their transcript annotations were selected from the original Gencode annotation file and placed into a separate file. To do this, a custom script called ‘selectGencodeAnnot.py’ was used to automatically search for and place all of the annotations for the 686 genes into a new abbreviated annotation (.gtf) file. The original Gencode annotation file is extremely large (765 Mb GTF file) and only 1.4% of the exon annotations pertained to the list of genes needed for analysis. This step further scaled down the complexity of subsequent analyses by reducing the number of annotated exons to analyze by 91.9% from 1,617,660 to 130,820.

*Collapsing redundant exons in the abbreviated GTF file.* Most genes express more than one splice isoform and the resulting transcripts often share multiple exons (e.g. many exons are constitutively expressed). This redundancy of exon annotations can make differential exon usage analysis difficult and inefficient. In addition, having multiple splice isoforms / different transcripts can lead to confusion in exon numbering (e.g. exon 4 of transcript-A is may have the same genomic location as exon 7 of transcript-B). In some cases, an exon of one transcript could contain shortened or extended nucleotide sequences 5’ or 3’ of an exon of an alternate isoform making analysis even more complex.

In order to reduce analysis complexity, we used the ‘dexseq_prepare_annotation.py’ script included in the DEXSeq software package (v1.28.1). This DEXSeq script takes the transcripts/splice isoforms belonging each gene and creates a single collapsed (‘meta’) annotation (Supplementary Figure S3A, red). This annotation has an independent and consistent numbering system that is ordered by chromosomal location. The new numbering system can also account for the existence of shorter and/or longer exons by dividing each coding region into discrete units referred to as exonic parts (for more detailed explanation see DEXSeq documentation). After collapsing the Gencode annotations (GTF) file for the 686 genes, the number of exon annotations to analyze was reduced by 80% from 130,820 exons to 26,513 unique exonic parts in the collapsed annotation (GFF) file. With the target gene and exonic part annotations ready, we could then determine exon usage.

*Turning qualitative cluster reads into quantitative full-length reads.* PacBio's real-time sequencing technology is known to produce many random insertion and deletion errors. Their IsoSeq method (bioinformatics processing pipeline) corrects for these errors by collapsing similar full-length reads into consensus (cluster) reads which are further improved through additional polishing using non-full length (NFL) reads. Although this approach greatly improves sequence quality and is useful for determining unique splice transcripts, the method reduces the quantitative nature of the data (e.g. a single cluster read could be produced from 1 full-length read or >5000 full-length reads). Fortunately, the IsoSeq method notes the number of full-length and NFL reads used to form each cluster read in the read names of the fastq file. PacBio states that this information should not be directly used for precise quantitative expression analysis (e.g. like Illumina RNA-Seq data). However, we sought to test if the data could be used quantitatively by validating our findings through RT-PCR. Therefore, in subsequent steps the 'exCovator.py' script, we extract and analyze only the full-length reads counts instead of cluster read counts. Its important to note that this can be muddled with collapsed reads, so do not use the ToFU pipeline.

*Finding unannotated exons from the data set.* Determining read coverage based on annotated exons alone neglects potentially novel/unannotated exons that may exist in our data. Long read technology has the opportunity to differentiate potential unannotated exons from sequencing and alignment artifacts seen in short-read data because assembly of transcripts is not required. In order to find unannotated exons, the ‘exCovator.py’ script uses the python HT-Seq library to: 1) sort all of the reads belonging to each input sample (BAM files) by the genes they map to, 2) iterate through each sample read and split into its constitutive exons using information provided in their CIGAR strings for every gene, 3) iterate through each exon of the read, 4) check to see if the exon’s genomic coordinates intersect with any annotated exons belonging to the same gene, 5) skip to the next read exon if there is a match, and 6) if the exon does not intersect with any annotated exons belonging to the gene, add the unannotated exon coordinates into the collapsed annotation file (Supplementary Figure S3B). Unannotated exons are first numbered, prefixed with a ‘u-’ (for differentiation), and sorted by starting coordinates before they are added alongside the list of annotated exons. Using this method, 1,218 unannotated exons were found in the three muscle samples, not present in the abbreviated Gencode annotation (GFF) file. This addition brings the total exonic parts to 27,731. Unannotated exons found depend on the sample (bam) and the annotation files given (e.g. the more comprehensive Gencode reference file will contain fewer unannotated exons than a RefSeq annotation file and samples from different tissues may express different transcript isoforms/exons).

*Normalizing exon coverage for cross sample comparisons.* Recalling the primary observations of the data (Figure 1), there was uneven exon coverage of genes within samples, especially 3’ end vs 5’ ends of very large genes due to the use of oligo-dT primers. In addition, there is sample-to-sample coverage variation for each gene from sequencing and library preparations. This variation makes it difficult to compare differential exon usage between samples. In the ‘exCovator.py’ script, this variation is accounted for by normalizing read coverage for each exonic part within each sample.

To normalize read coverage, a fraction (ratio similar to percent spliced in or PSI) is calculated from the total population of reads mapping to each exonic part (Supplementary

Figure S3C). The denominator for the fraction is a total count of reads that have genomic coordinates intersecting the exonic part's coordinates. This value will be referred to as the 'total reads' for the exonic part. Reads that have intersecting coordinates with the exonic part (total reads) may or may not have a sequence match with the exonic part. The numerator of the fraction is the subpopulation of total reads that contain an exact sequence match to the annotated exonic part in question. To check if any of the reads in the population of total reads have a sequence match to the annotated exonic part, a check is performed on each read one by one. The script breaks each read into its constituent exons using the exon boundary information contained in its CIGAR string. Next, each exon of that read is checked to see if its coordinates intersect that of the annotated exonic part. If it does, it is counted as a 'match read'. This allows for the analysis to tolerate reads containing some inaccurate base calls since we are analyzing the data at the exon level. However, the analysis can produce artifacts if the read contains indels at an exon junction, is mismapped or if the sequencing data is contaminated with DNA (reads containing introns). Therefore, some manual inspection of reads in IGV is required.

Using this method, the total count, and match count for all 27,731 annotated and unannotated exonic parts were calculated. Finally, the population of match reads is divided by the total reads to generate a normalized ratio or PSI for each exonic part (PSI = match / total = ratio * 100). This ratio can be used to compare differential exon usage within and across different samples. However, this will only determine coverage for all annotated (known) exonic parts. After the match reads, total reads, and ratios were calculated for each exon in all three of the samples, the data was output into a CSV file. However, this large sum of exonic parts and the resulting data table was going to be challenging to analyze manually.

*Visualizing differential exon usage between tissues.* The CSV data file with all of the ratios contained too many exons to compare between samples by eye. Therefore, an intuitive way to visualize differential exon usage between samples would be useful for analysis. The ‘exCovator.py’ script generates a PDF file containing coverage plots for each gene. Each gene plot is composed of two types of graphs that share a common x-axis (Figure 4A, 5A and 6A). This x-axis displays all of the exonic parts (annotated and unannotated) for the gene ordered by chromosomal location. The top graph represents each sample as a colored line. Its y-axis displays the coverage ratio (full-length match/total read counts) for each exonic part. A ratio of 1 represents 100% of the total reads of the exon are all sequence matches. However, a ratio of zero is more ambiguous; a zero can represent no match reads or no total reads. In addition, a low denominator (total reads) can make ratios unreliable and/or provide less confidence in the data (e.g. a 0.5 ratio could be match/total of 300/600 or 2/4). To remove this ambiguity, we added a stacked bar graph below the line graph that shows the total reads for each sample. This way the user can assess both the total read count and differential exon usage in one glance to determine if the exon is worth analyzing further or rule out as an artifact. Once it was possible to visualize the data and find differentially used exonic parts, it would still be useful to filter and and obtain additional information about the exonic parts from the original CSV data file.

*Filter for differentially used exons between samples*. Most exons of a gene are used in all splice isoforms across multiple tissues (constitutively expressed). However, some exons are not used at all in splice isoforms of specific tissues (e.g. muscle vs brain). Both cases are examples of exons that will not be differentially usage between samples; the ratios remain constant in all samples and become additional noise in the analysis (Supplementary Figure S3D, black percentages). In order to focus only on exons that are differentially used between samples (Supplementary Figure S3D, red percentages), the ‘FilterDiffUsedExons.py’ script was created to filter out exons that did not change between samples. A percent change cutoff was used to change the filter stringency. We chose to apply a filter cutoff of at least a 10% difference in exon coverage ratio between all samples. This is a lenient cutoff, allowing for most differentially used exons to be included. However, some exons that have very few reads may fall below 10%

differentially used threshold. These rare and potentially novel exons will be filtered out of this analysis, but we were interested in capturing the most robust changes in this first analysis (we use exPhaser for more sensitive analysis). The script yielded 2,631 exonic parts in 433 genes that were at least 10% differentially used between the three muscle samples. This filtering step reduced the amount of exonic parts to analyze by 90% (27,731 to 2,631), a more manageable number but still an extremely tedious task to perform manually.

*Selecting exonic parts by gene function.* Manually analyzing 2,631 exonic parts in 443 genes would have been a labor-intensive task. In addition, many of these exons may have little to do with the focus of our study: very large muscle structural genes. Therefore, additional information about the 443 genes would help determine their relevance to muscle such as their full names (rather than just gene symbols) and function. The NCBI gene database contains comprehensive summaries for each gene and an application-programming interface (API) for users to automate queries and obtain data. In order to obtain additional gene information from NCBI, we created a script called 'selectGeneSummaries.py' that uses two python libraries: MyGene and BioPython. The goal was to convert the 443 gene symbols into NCBI gene ids. These ids could then be used in a separate query to obtain the gene summaries using NCBI's batch Entrez. The BioPython library makes automating the queries with NCBI's databases much simpler. However, doing two large queries: 1) for 443 gene ids and 2) for their summaries could reach NCBI's user limit for database queries. Going over the limit can result in a permanent IP address ban. Therefore, the MyGene library was used to rapidly query for NCBI gene ids which has no limitations on the number of queries. Then BioPython was used to do an NCBI Batch query with the gene ids to download all the gene summaries for the 443 genes containing differentially used exons into a CSV file. Finally, we searched through the gene summaries and found 43 for genes related to muscle, 15 of which were structurally related. By automating the retrieval of the full gene names and their functions, we were able to markedly reduce the number of genes of interest by 97% for analysis. We chose Nrap, Nebulin and Titin for further analysis.

*Manual (bioinformatic) verification of differentially used exonic parts.* The `exCovator.py` script finds unannotated exons by looking for sequences in the samples that are not part of the annotation (GFF) file. This method can lead to potential novel discoveries, but also lead to false positives due to sequencing indels, mapping errors, or PCR artifacts during cDNA generation. In order to differentiate likely novel candidates from potential false positives, we manually cross checked the exonic part data using a combination of observations from the IGV browser, the coverage graphs, and the coverage ratios calculated from the `exCovator.py` script. Using these tools, exon candidates were selected using the following guidelines: 1) each candidate should have a reasonable amount of reads in at least one of the samples (e.g. >= 30 total full-length read coverage in at least one of the samples), 2) there should be a marked difference in coverage ratios between at least two of the three samples (e.g. >= 20%), 3) multiple reads containing the sequence matched exon should have the same genomic interval (start and end coordinates), and 4) a majority of the reads should not contain insertion or deletion errors at the splice sites/exon boundaries and 5) if it is an unannotated exon, it should be conserved in other species.

*Scripts for this analysis pipeline can be found in the Github repository.*
[www.github.com/puapinyoying/isoseq_manuscript_resources/s2_exCOVator_analysis_pipeline](www.github.com/puapinyoying/isoseq_manuscript_resources/s2_exCOVator_analysis_pipeline)

**exPhaser transcript structure analysis pipeline** (supplementary data – exPhaser implementation pdf)

The exCovator analysis pipeline is exon-based so it is helpful for identifying unannotated exons and quantifying differential usage of individual exons between tissues. However, it cannot determine exon phasing. The goal of the exPhaser analysis pipeline is to determine exon phasing pattern of multiple key (ideally isoform defining) exons within a transcript and identify and quantify the total transcripts in the given samples. (See ‘exPhaser implementation’ PDF slides in supplemental data for examples and Figures 6 and 7).

*Generate Boolean reference table to count all possible splice patterns.* First, the script takes the user list (BED) of key exons to phase and calculates the total possible splice patterns using this formula: 2^number of exons. Next, the total possible patterns starting from zero (e.g. 1-8 patterns would be 0-7 patterns), were converted to binary numbers (e.g. 0 = 000, 1=001, 7=111, etc) and split into a table and converted into a Boolean (True/False) table containing presence and absence of all exon combinations.

*Calculate interval range to select reads and annotations for analysis.* In order to normalize the data for comparisons between each sample, we need to calculate a denominator of total reads (that span the range of exons, matching the sequence or not). Not all the reads that map to a gene of interest will span the list of key exons provided for phasing analysis. In order to filter out reads that do not cover the exons of interest, we calculate an interval range variable which is the start coordinate of the first exon and the end coordinate of the last exon given by the user (pre-sorted by genomic coordinates). The reads that *contain* the interval range are selected for further analysis. The same is done for the given Gencode annotation (gtf) file to select annotations that contain the interval range.

*Annotate and count each splice pattern in the Boolean reference table.* Using HT-Seq, each annotation and read that contained the interval range was checked for its splicing pattern. If the splice pattern of the user provided exons matched one or more annotations, then they are annotated in the pattern's row in Boolean table. For each sample read, if the splice pattern found in the cigar string matched the Boolean table, then we would start a running count for it.

*Filter and normalize the count data.* After the table is filled with count and annotation data, it is filtered to remove splice patterns that have a zero read count or if it is not annotated. Then the data is normalized by dividing the number of reads for each transcript structure/splice pattern by the total full-length read count of all reads that contain the interval range. Finally, the data table is exported for manual analysis.

*Scripts for this analysis pipeline can be found in the Github repository.*

www.github.com/puapinyoying/isoseq_manuscript_resources/s3_exPhaser_analysis_pipeline
