## Supplementary material for "A new long-read RNA-seq analysis approach identifies and quantifies novel transcripts of very large genes": exPhaser_implementation

Program for Determining phasing of isoform defining exons

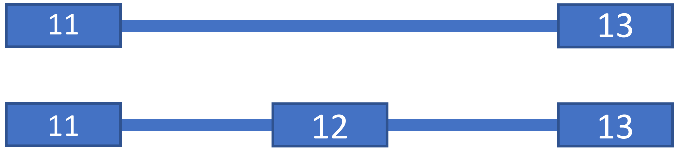


Brian Uapinyoying

Hoffman lab

8/6/2019

### Introduction

- PacBio's long-read isoform sequencing data spans multiple exons (10kb+)
- This allows for phasing of exons when analyzing RNA-Seq data
- But some transcripts are too long > 10 kb (e.g. Ttn at 106 kb)
- We successfully used an exon-based approach (ExCOVator) and internal priming to determine differential usage between tissues in ultra long transcripts
- However, without phasing data we cannot determine the splicing pattern of these differentially used exons within individual transcripts from each tissue

### Why is phasing of exons important?

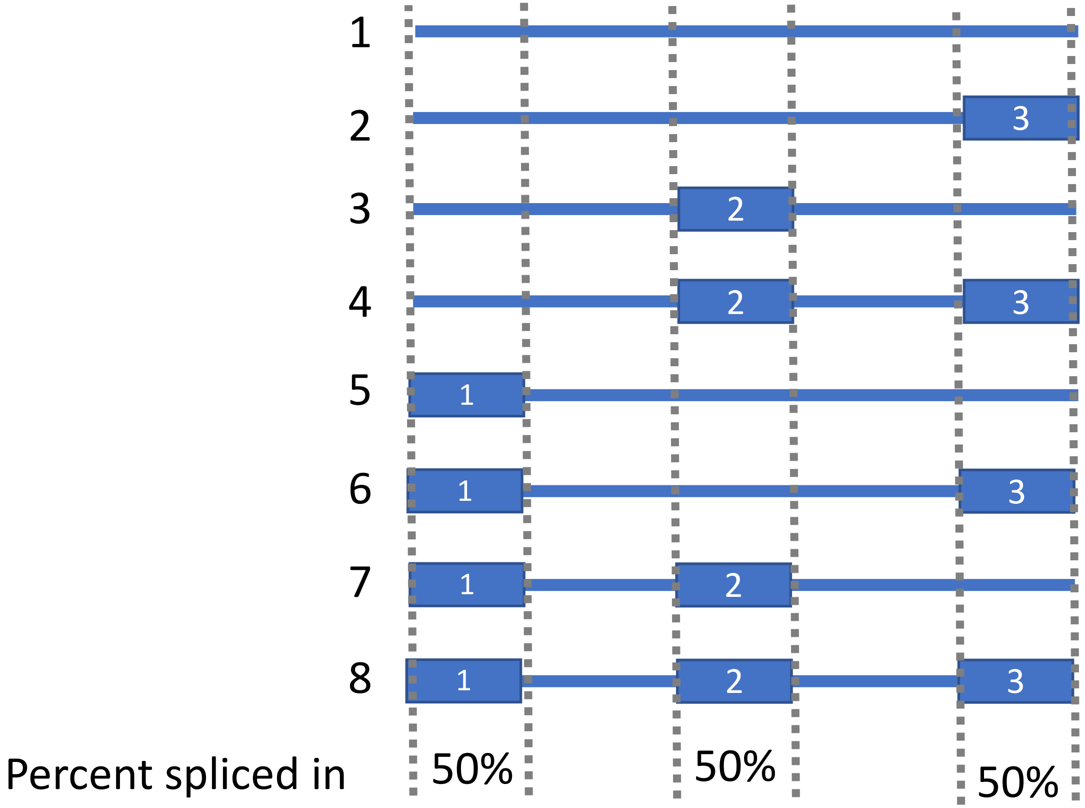


- Without phasing data,  all 3 exons (independently) look like they are similarly expressed when they could be part of different transcript isoforms

### General Implementation

- Use the python library HT-Seq
  - https://www.ncbi.nlm.nih.gov/pubmed/25260700
- Select  key exons / isoform defining cassette exons
  - input into script as bed file (genomic coordinates)
- Loop through each IsoSeq read from sample bam files
  - Determine if the read is long enough to cover all key input exons
    - Filter out reads that are too short
  - Check if exons in the read match any of the key input exons & note splice pattern
  - Tally up all the reads for each splice pattern and output as data table/file

### What are all the possible splicing patterns with a given number of exons?

- Formula = $2^{(\#exons)}$
- For 3 exons, $2^3$ = 8 total splicing possibilities or 8 rows of information
- Each row (i) can be seen as binary representation of [row number – 1]

Number of rows = $2^{(\#exons)}$

(row# – 1)

| Row # | Binary |
|---|---|
| 0 | 000 |
| 1 | 001 |
| 2 | 010 |
| 3 | 011 |
| 4 | 100 |
| 5 | 101 |
| 6 | 110 |
| 7 | 111 |

Split into Binary table

| Exon 1 | Exon 2 | Exon 3 |
|---|---|---|
| 0 | 0 | 0 |
| 0 | 0 | 1 |
| 0 | 1 | 0 |
| 0 | 1 | 1 |
| 1 | 0 | 0 |
| 1 | 0 | 1 |
| 1 | 1 | 0 |
| 1 | 1 | 1 |

Convert to Boolean values
(Exon Spliced in? True/False)

| Pattern | Exon 1 | Exon 2 | Exon 3 |
|---|---|---|---|
| 1 | False | False | False |
| 2 | False | False | True |
| 3 | False | True | False |
| 4 | False | True | True |
| 5 | True | False | False |
| 6 | True | False | True |
| 7 | True | True | False |
| 8 | True | True | True |

### Not all possible patterns will exist due to constitutively expressed exons

**Hypothetical Example**: NRAP Soleus

Likely only a subset of the total possibilities will be seen in the data, the rest may be artifacts

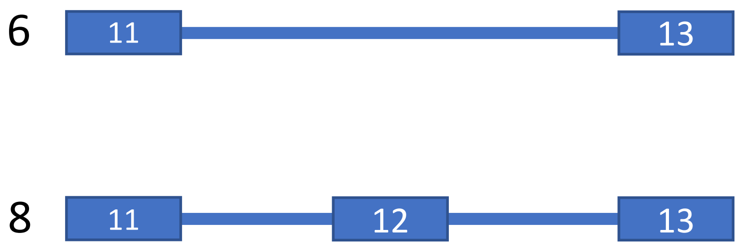


| Pattern | Exon 11 | Exon 12 | Exon 13 | FL_count | % |
|---|---|---|---|---|---|
| 1 | False | False | False | 2 | 0 |
| 2 | False | False | True | 5 | 0 |
| 3 | False | True | False | 0 | 0 |
| 4 | False | True | True | 1 | 0 |
| 5 | True | False | False | 0 | 0 |
| 6 | True | False | True | 1500 | 30 |
| 7 | True | True | False | 0 | |
| 8 | True | True | True | 3500 | 70 |

### Technically, we also do not have to limit ourselves to adjacent (neighboring) exons as long as the reads are long enough to contain all exons

- If we know which exons are variably spliced (output from my exCOVator script), we can strategically select these key exons for phase analysis
- Then determine presence/absence of input exons relative to each other within each read
- This same approach can be done for transcript annotations!
- However, the longer the distance (interval range) between the first and last key exon, the fewer reads will contain all key exons

| Pattern | Exon 9 | Exon 17 | Exon 47 |
| --- | --- | --- | --- |
| 1 | False | False | False |
| 2 | False | False | True |
| 3 | False | True | False |
| 4 | False | True | True |
| 5 | True | False | False |
| 6 | True | False | True |
| 7 | True | True | False |
| 8 | True | True | True |

### Only reads and transcript annotations that **contain** the bed/exon interval range are selected for determining and quantifying splicing patterns

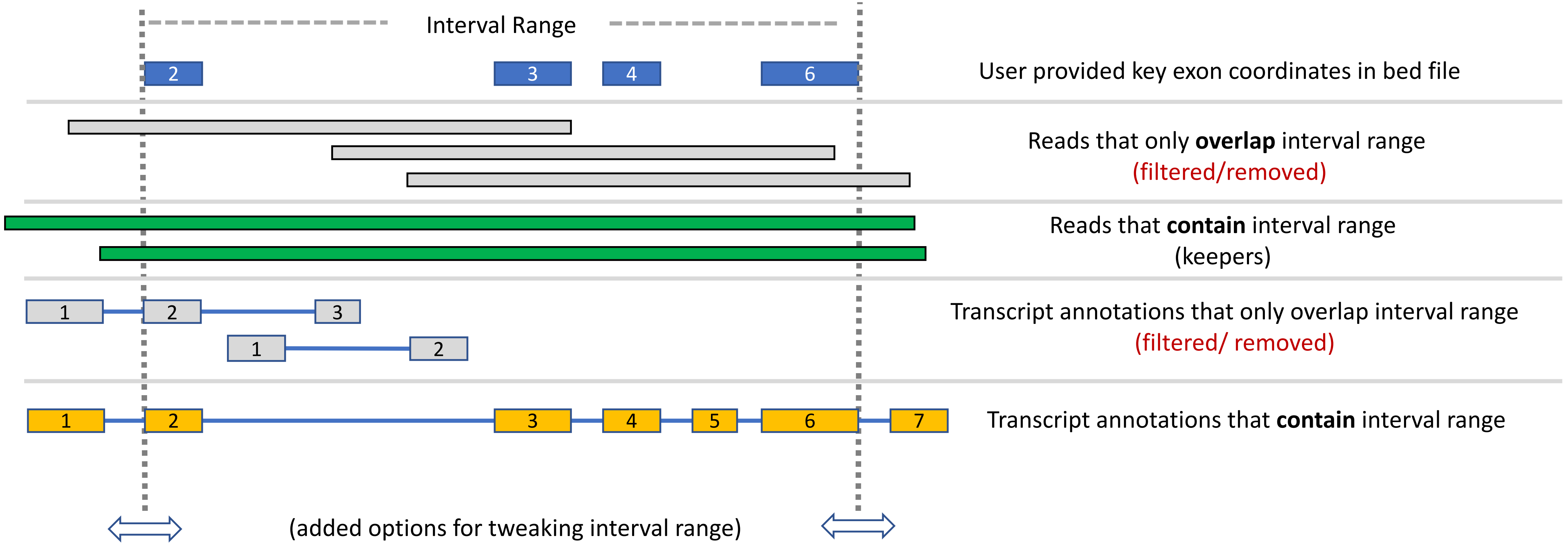


- Interval range is calculated using the start site of the lowest coordinate exon and the end site of the highest coordinate exon
- Therefore, the farther the first and last exons are the fewer reads will contain the interval range
- Reads that overlap vs contain the interval range is output to a data file for the user

### We can also use this same approach to determine which pattern is associated with which annotated transcripts

| Isoform | Exon 9 | Exon 17 | Exon 47 | Annotation |
|---|---|---|---|---|
| 1 | False | False | False | ? |
| 2 | False | False | True | NM_XX3 |
| 3 | False | True | False | ? |
| 4 | False | True | True | ? |
| 5 | True | False | False | ? |
| 6 | True | False | True | NM_XX2 |
| 7 | True | True | False | ? |
| 8 | True | True | True | NM_XX1, NM_XX4 |

Most patterns won't be annotated or exist

Some patterns may match multiple transcript annotations

### Finally, we can combine all three pieces of data (pattern, annotation, read count) and filter out patterns with no reads in any sample and with no annotation found

All patterns

| Isoform | Exon 9 | Exon 17 | Exon 47 | Annotation | Cardiac Read count | EDL Read count | Soleus Read count |
|---|---|---|---|---|---|---|---|
| 1 | False | False | False | ? | 0 | 0 | 0 |
| 2 | False | False | True | NM_XX3 | 100 | 700 | 300 |
| 3 | False | True | False | ? | 2 | 0 | 0 |
| 4 | False | True | True | ? | 0 | 250 | 0 |
| 5 | True | False | False | ? | 0 | 0 | 0 |
| 6 | True | False | True | NM_XX2 | 30 | 0 | 900 |
| 7 | True | True | False | ? | 0 | 0 | 0 |
| 8 | True | True | True | NM_XX1, NM_XX4 | 500 | 0 | 40 |

Selected patterns

| Isoform | Exon 9 | Exon 17 | Exon 47 | Annotation | Cardiac Read count | EDL Read count | Soleus Read count |
|---|---|---|---|---|---|---|---|
| 2 | False | False | True | NM_XX3 | 100 | 700 | 300 |
| 3 | False | True | False | ? | 2 | 0 | 0 |
| 4 | False | True | True | ? | 0 | 250 | 0 |
| 6 | True | False | True | NM_XX2 | 30 | 0 | 900 |
| 8 | True | True | True | NM_XX1, NM_XX4 | 500 | 0 | 40 |

Artifact or very rare transcript?

Potential novel isoforms

Not enough data to distinguish the two isoforms

### **Pilot Analysis**: Nrap transcript expression between Cardiac, EDL and Soleus using Gencode mm10 annotations

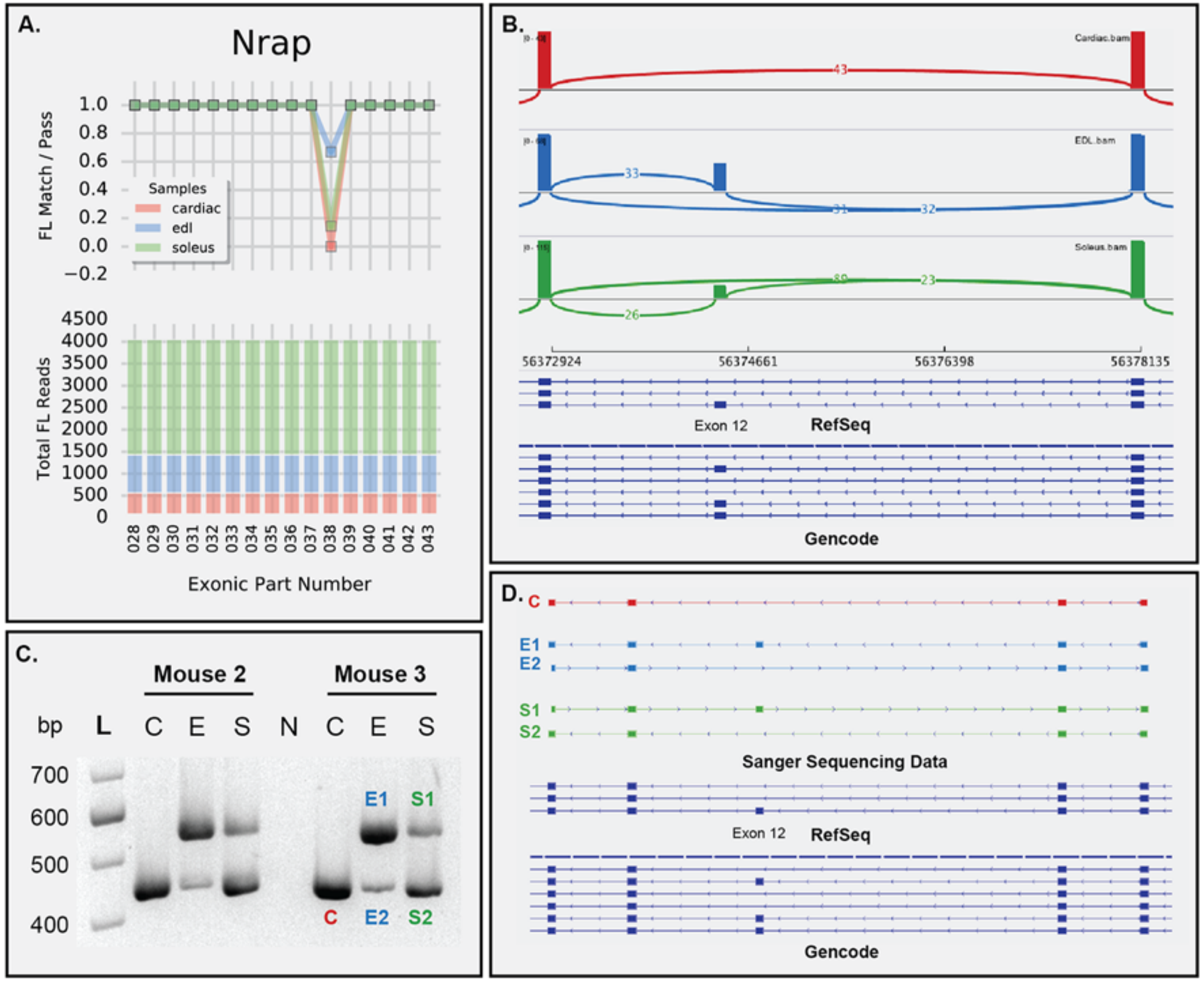


- Exon 12 was differentially spliced between tissues
- Are there other exons we didn't initially detect?

### Difficult to tell based on eyeballing reads (IGV)

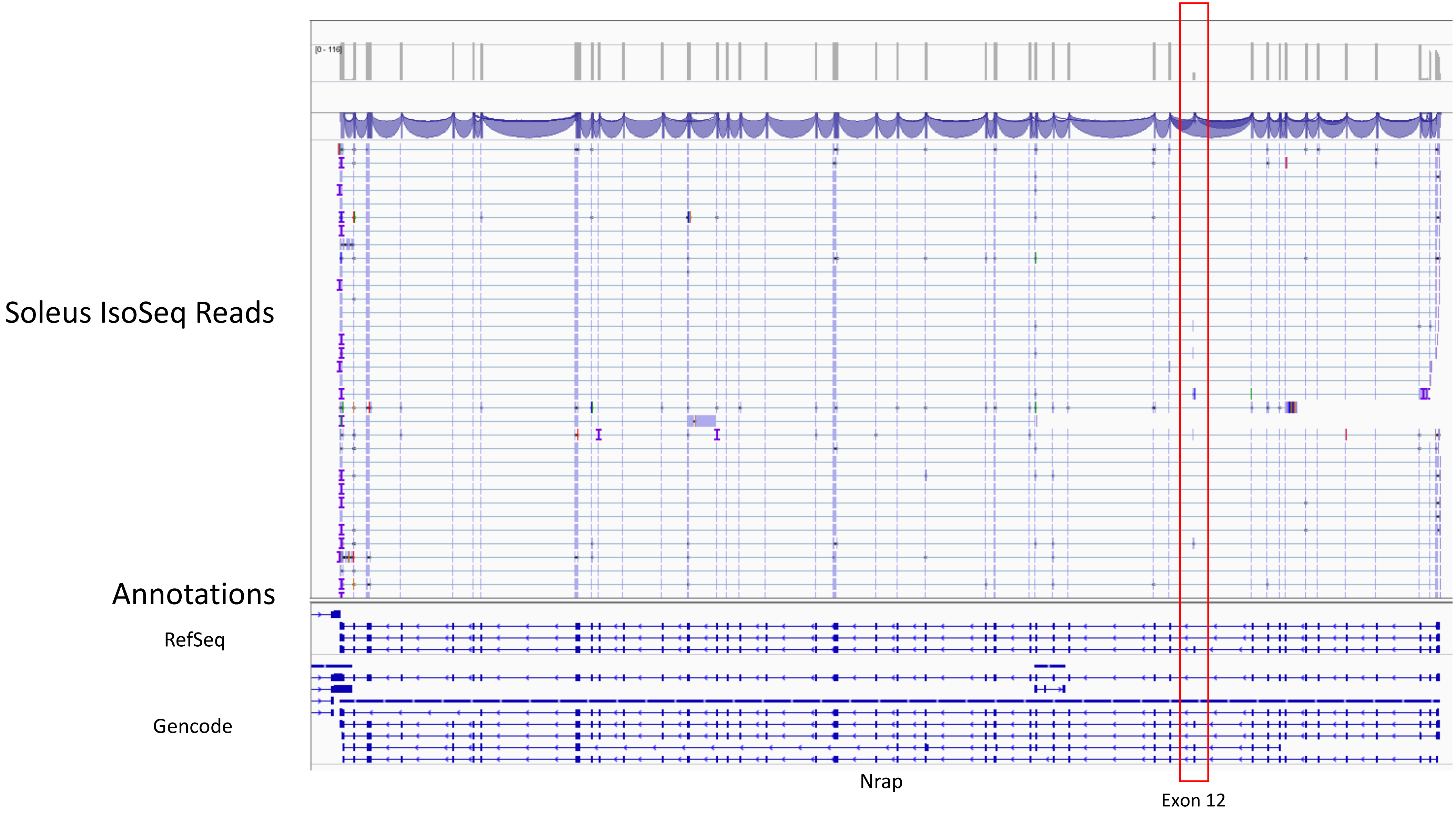

### **Example Analysis:** Nrap transcript expression between Cardiac, EDL and Soleus using Gencode mm10 annotations

- Exons 2, 12, 37, 38, 39 and 40 are cassette/isoform defining exons (6 total)
    - Omitting ENSMUST00000169099.7 which is short and has been labed as 'non-sense mediated decay'
- Added a 6 constitutively expressed adjacent/neighboring exons as anchors for analysis, exons 41, 36, 13, 11, 3, 1
    - Not required for phasing to work, but adds some context and helps validate findings

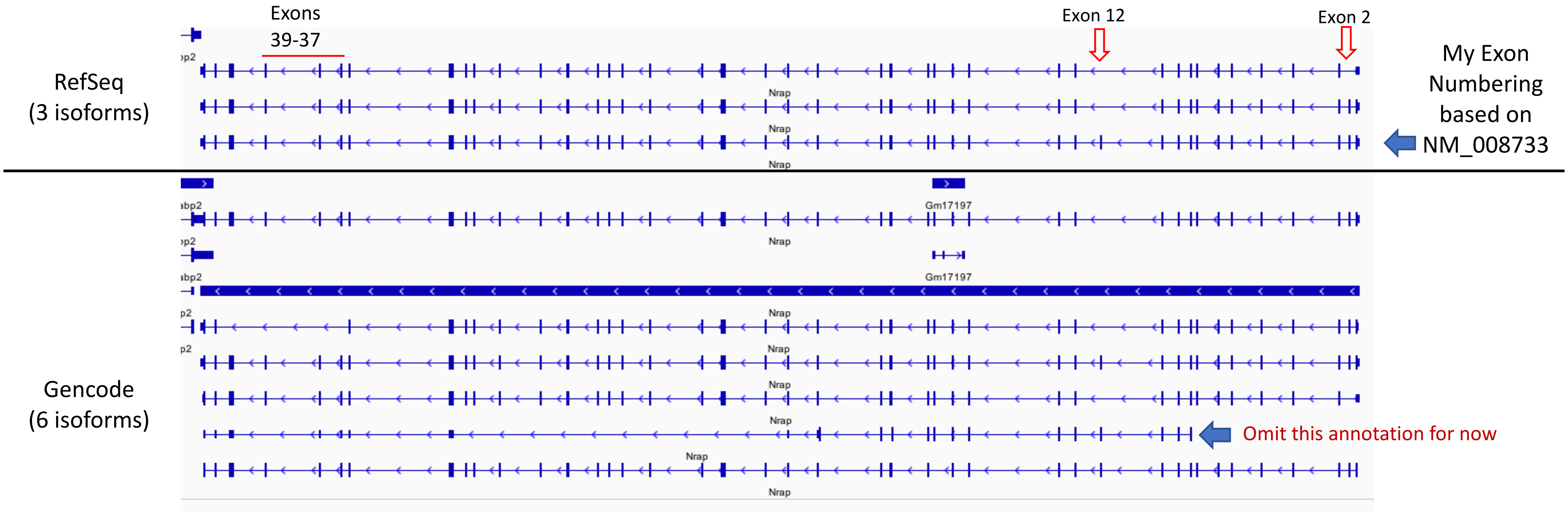

### Input bed file (genomic coordinate file separated by tabs) for Nrap

| Chrom | Start | End | Name | Score | Strand |
|---|---|---|---|---|---|
| chr19 | 56320884 | 56321016 | exon41 | 0 | - |
| chr19 | 56321722 | 56322034 | exon40 | 0 | - |
| chr19 | 56323887 | 56323995 | exon39 | 0 | - |
| chr19 | 56327194 | 56327299 | exon38 | 0 | - |
| chr19 | 56328500 | 56328599 | exon37 | 0 | - |
| chr19 | 56328995 | 56329100 | exon36 | 0 | - |
| chr19 | 56372806 | 56372914 | exon13 | 0 | - |
| chr19 | 56374363 | 56374468 | exon12 | 0 | - |
| chr19 | 56378054 | 56378171 | exon11 | 0 | - |
| chr19 | 56388748 | 56388836 | exon3 | 0 | - |
| chr19 | 56389389 | 56389484 | exon2 | 0 | - |
| chr19 | 56389805 | 56390038 | exon1 | 0 | - |

Red exon names are isoform defining exons

- Original coordinates were extracted from UCSC Table Browser (RefSeq mm10 for NRAP)
- Modified name column to make it easier for me to remember the exons
- Input to script also requires sample bam files and gencode annotation (gtf) file selected for the target gene
  - cardiac.bam, edl.bam, soleus.bam, nrap.gtf, nrap_exons.bed

### Basic statistics on reads that contained interval range (keepers used in phasing analysis) vs total reads overlapping interval range (all reads)

| | Reads that Contain Interval range | | Reads that overlap interval range | | Percent contained / overlap | |
|---|---|---|---|---|---|---|
| **Sample** | **Cluster count** | **Full-length count** | **Cluster count** | **Full-length count** | **Cluster count** | **Full-length count** |
| cardiac | 12 | **399** | 43 | **567** | 27.9% | **70.4%** |
| edl | 12 | **483** | 68 | **863** | 17.6% | **56.0%** |
| soleus | 32 | **1771** | 116 | **2622** | 27.6% | **67.5%** |

- **Cluster count** = Pacbio's cluster/consensus read that is output from its ICE algorithm. Basically grouping multiple full-length reads by similarity and using it to increase sequence quality by forming in a single unique polished read

- **Full-length count** = number of full-length reads extracted from information in the cluster read (more accurate)

### **Results** for Nrap show that we are able to determine transcript level information from phasing of key exons in the annotation and read data

| Exon 41 | Exon 40c | Exon 39c | Exon 38c | Exon 37c | Exon 36 | Exon 13 | Exon 12c | Exon 11 | Exon 3 | Exon 2c | Exon 1 | transcript_ids | Cardiac flCount | Cardiac pct | Edl flCount | Edl pct | Soleus flCount | Soleus pct |
|---|---|---|---|---|---|---|---|---|---|---|---|---|---|---|---|---|---|---|
| TRUE | FALSE | FALSE | FALSE | FALSE | TRUE | TRUE | FALSE | TRUE | TRUE | TRUE | TRUE | ENSMUST00000167239.7 | 0 | 0% | 0 | 0% | 0 | 0% |
| TRUE | TRUE | TRUE | TRUE | TRUE | FALSE | TRUE | FALSE | TRUE | TRUE | TRUE | TRUE | ? | 0 | 0% | 0 | 0% | 4 | 0% |
| TRUE | TRUE | TRUE | TRUE | TRUE | TRUE | TRUE | FALSE | TRUE | TRUE | FALSE | TRUE | ENSMUST00000095947.10 | 14 | **4%** | 0 | 0% | 1 | 0% |
| TRUE | TRUE | TRUE | TRUE | TRUE | TRUE | TRUE | FALSE | TRUE | TRUE | TRUE | TRUE | ENSMUST00000040711.14 | 385 | **96%** | 153 | **32%** | 1553 | **88%** |
| TRUE | TRUE | TRUE | TRUE | TRUE | TRUE | TRUE | TRUE | TRUE | TRUE | FALSE | TRUE | ? | 0 | 0% | 0 | 0% | 1 | 0% |
| TRUE | TRUE | TRUE | TRUE | TRUE | TRUE | TRUE | TRUE | TRUE | TRUE | TRUE | TRUE | ENSMUST00000073536.12, ENSMUST00000166203.1 | 0 | 0% | 330 | **68%** | 212 | **12%** |
| | | | | | | | | | | | | **total** | **399** | | **483** | | **1771** | |

- ENSMUST00000073536.12, and ENSMUST00000166203.1 are almost identical except for lengths of 3’ and 5’ UTR
  - Indistinguishable with current Interval range
- Some reads in cardiac for ENSMUST00000095947.10 wasn’t seen before
  - Has low coverage (14 reads). Could be rare transcript.
- Transcript level analysis shows similar ratios of differential expression as exon 12 RT-PCR
- The two unannotated matches (?) were checked in IGV to be from artifact reads

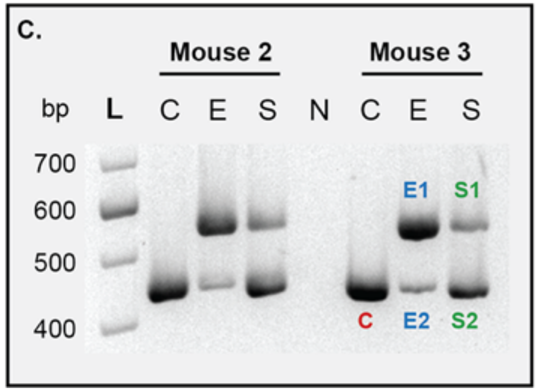


RT-PCR

With Exon 12

Without Exon 12
